# The chloroplast CLPD chaperone: consequences of under- and overexpression, interaction with the CLP protease core, and candidate substrates

**DOI:** 10.64898/2026.05.10.723991

**Authors:** Marissa Y. Annis, Pratyush Routray, Nazmul H. Bhuiyan, Bingjian Yuan, Klaas J. van Wijk

## Abstract

Expression of the chloroplast AAA+ chaperone *CLPD* gene increases during senescence and drought, but its function in chloroplast proteostasis is poorly understood. This study provides a comprehensive analysis of Arabidopsis CLPD protein accumulation across development from early seedlings to senescence, and compares results to its homologs CLPC1,2, as well as CLPB3 and cpHSP90. The developmental consequences of complete loss of CLPD expression (*clpd-1*), as well as overexpression of functional CLPD or CLPD impaired in ATP hydrolysis (CLPD-TRAP), were determined in Arabidopsis. *clpd-1* has accelerated seedling development while functional CLPD overexpression lines, but not CLPD-TRAP, have delayed development. To determine if CLPD is a *bona fide* CLP chaperone associating with the CLPPRT protease and to identify *in vivo* candidate substrates, we employed the CLPD-TRAP line during vegetative and flowering (senescent) growth stages. Affinity purification of CLPD-TRAP followed by mass spectrometry showed high enrichment of all nine subunits of the CLP protease, suggesting that CLPD plays a role in substrate delivery to the CLP protease. CLPC1,2 were also highly enriched in the CLPD-TRAP interactome, and together with prior information, suggesting hetero-oligomerization of the three chaperones. Nine chloroplast candidate substrates were identified in the CLPD-interactomes, including: FHY2 involved in riboflavin synthesis, THI1 and THIC involved in thiamin metabolism, and four proteins of unknown function. Several of these were previously identified as potential CLP substrates based on CLPC1 trapping and comparative proteomics of c*lp* mutants. Together, this suggests that CLPD acts in substrate selection within a heteromeric CLPC-CLPD hexamer, likely with unique contributions.

## Introduction

ATP-dependent CLP proteases are present in bacteria, mitochondria and plastids, and regulate accumulation levels of a broad range of substrates (Sauer and Baker, 2011; Alexopoulos et al., 2012; Liu et al., 2014; Nishimura and van Wijk, 2015; Nishimura et al., 2017). The CLP machinery components greatly diversified during evolution from prokaryotic progenitors, with plant chloroplasts containing the most complex CLP system. The protease core of the chloroplast CLP system in *Arabidopsis thaliana* is a tetradecameric assembly of five proteolytically-active subunits (CLPP1, CLPP3–6) and four proteolytically-inactive proteins (CLPR1–4) in a fixed stoichiometry of one to three copies each (Olinares et al., 2011). The Arabidopsis chloroplast CLP system also features two stabilizing/activating factors (CLPT1,2), three Class I AAA+ chaperones (CLPC1, CLPC2, and CLPD), the adaptor CLPS1, and the plant-specific co-adaptor CLPF (Peltier et al., 2004; Nishimura et al., 2013; Kim et al., 2015; Nishimura et al., 2015).

Arabidopsis CLPC1 and CLPC2 share around 90% sequence identity and more than 70% sequence identity with cyanobacterial CLPC, whereas CLPD is only found in higher plants and shows only 45% identity to CLPC proteins in plastids and cyanobacteria. CLPC1,2 and CLPD contain two AAA domains that hydrolyze ATP to power substrate unfolding (Fig. 1). All three chaperones also possess a conserved short hydrophobic motif within the P-loop near the C-terminus and the IGF/L motif that is essential for association of chaperones with the CLPP protease core in bacteria (Kim et al., 2001; Singh et al., 2001; Kress et al., 2009) (Fig. 1). Located just a few residues immediately downstream of the IGF loop in CLPC chaperones is the R-motif, a short eight amino acid sequence rich in basic residues that confers specific interaction between CLPC and the R ring of the CLP core in cyanobacteria (CLPP1, CLPR1-4 in Arabidopsis). The R-motif is also conserved in plastid CLPC1,2 (Tryggvesson et al., 2012), but is only partly conserved in CLPD (Fig. 1). However, it is unknown if the R motif is important in angiosperm CLPC chaperones.

**Figure 1.**
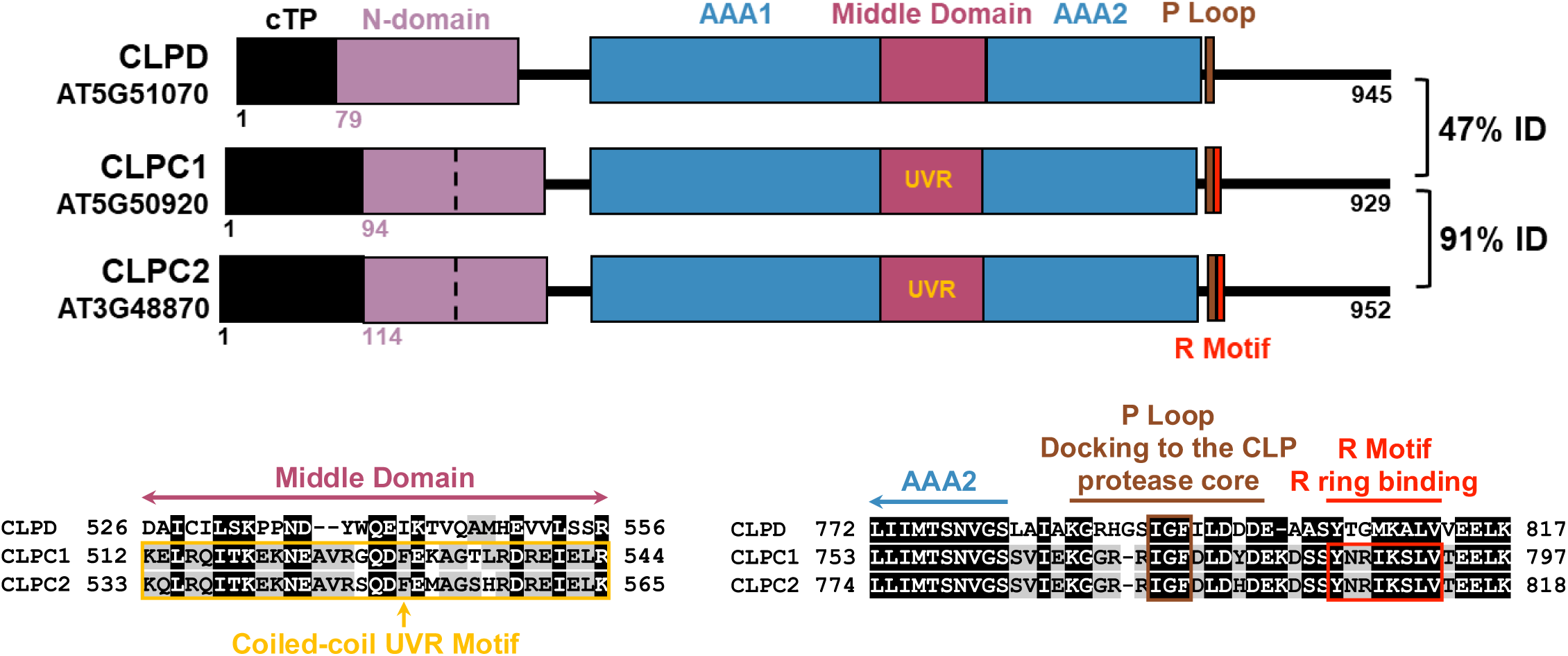
Chloroplast chaperones of the CLP protease system in Arabidopsis. Domain map of chloroplast CLP chaperones based on primary amino acid sequence. Percent sequence identity (% ID) was calculated from a MUSCLE alignment (Madeira et al., 2022) of the amino acid sequence of the mature proteins without chloroplast transit peptides. Assignment of N-termini of mature CLPC1,2 and CLPD are based on sequence conservation of plant CLP chaperone homologs and MSMS-based protein data available in PPDB and PeptideAtlas. Dashed lines signify the two-fold symmetry of the N-domain, which is broken in CLPD. UVR denotes the presence of a coiled-coil UVR motif within the Middle domain (see (Annis et al., 2024). Also shown (below) are regions of a sequence alignment of Arabidopsis CLPD, CLPC1, and CLPC2 generated with MUSCLE (Madeira et al., 2022). Residues are colored according to similarity using the BLOSUM62 scoring matrix. Shown on the left is the CLPD sequence (D526-R556) aligning to the conserved UVR motif (yellow) sequences in CLPC1 (K512-R544) and CLPC2 (K533-K565). These regions in CLPC1 and CLPC2 are predicted to form the canonical coiled-coil structure of UVR motifs, which regulate ClpC chaperone activity and oligomerization in bacteria (Carroni et al., 2017). Shown on the right is the alignment of the regions of CLPD, CLPC1, and CLPC2 that contain the IGF loop (brown) and the R Motif (red). The IGF tripeptide in the P loop is required for docking of the chaperone to the CLPP protease core in bacteria (Weibezahn et al., 2004). The R motif sequence is required for docking to the heteromeric CLPPR protease core in cyanobacteria and is lacking in CLPD (Tryggvesson et al., 2012).

*In vitro*, CLPC1,2 and CLPD each form homodimers but can form homohexamers in the presence of magnesium and ATP (Rosano et al., 2011; Jagdev et al., 2021). Hexamers are the active oligomeric state of CLP chaperones in bacteria, and chaperone hexamers from functional complexes with the CLP protease core complex for substrate unfolding and degradation (Kirstein et al., 2006; Kar et al., 2008). The CLPD Middle domain lacks the coiled-coil UVR motif conserved in CLPC1,2 and bacterial CLPC chaperones (Fig. 1). In bacterial CLPC proteins, this UVR motif plays a role in regulating chaperone oligomerization and regulating rates of ATPase activity (Carroni et al., 2017; Annis et al., 2024). The ATP hydrolysis rates of recombinant CLPC2 *in vitro* are comparable to those of CLPA in *E. coli* or CLPC in cyanobacteria (Rosano et al., 2011). In contrast, recombinant CLPD is kinetically slower compared to recombinant CLPC2 with a much higher Km value for ATP, but mixing with CLPT stimulates ATP hydrolysis rates (Rosano et al., 2011). Likewise, *in vitro* renaturation activity of a heat-aggregated model substrate is lower for CLPD than CLPC2 (Rosano et al., 2011). It has been shown that CLPC2 and CLPD are both capable of interacting with the N-terminal presequence (cTP) of the chloroplast pea protein ferredoxin *in vitro*, but the physiological significance of this is not clear (Bruch et al., 2012).

The specific *in vivo* substrate preferences of CLPC1,2 and CLPD are not known. The N-domains of CLP chaperones, as well as their middle domains, are important in binding of substates and adaptors (Olivares et al., 2018; Mabanglo and Houry, 2022). The N-domain of chloroplast CLPC1 is involved in binding of the adaptors CLPS1 and CLPF, as well as the substrate glutamate t-RNA reductase, whereas CLPF also interacts with the middle domain of CLPC1 (Nishimura et al., 2013; Nishimura et al., 2015). The structures of the Arabidopsis CLPC1 and CLPD N-domain have been resolved at atomic resolution by X-ray crystallography (Mohapatra et al., 2017) which showed that the CLPD N-domain has divergent structural features from CLPC and other CLPC/A homologs, suggesting a possible differences in substrate recognition (Mohapatra et al., 2017) (Fig. 1).

Arabidopsis CLPC1,2 mRNA and protein levels are highest in the first true leaves and the youngest leaves within a rosette and decline as the leaves mature (Nakabayashi et al., 1999; Nishimura et al., 2013; Sjogren et al., 2014). In contrast, CLPD levels generally increase in older leaves (Nakabayashi et al., 1999; Sjogren et al., 2014) and mRNA levels are strongly induced during leaf senescence (Nakashima et al., 1997; Olinares et al., 2011). However, there have been conflicting reports over whether CLPD protein increases during senescence (Weaver et al., 1999; Sjogren et al., 2014); therefore, a more comprehensive analysis of CLPD mRNA and protein levels throughout development is needed. *CLPD* gene expression is under the control of three NAC family transcriptional activators and a zinc finger homeodomain transcriptional activator, all of which are induced during drought, high salinity and abscisic acid (Simpson et al., 2003; Tran et al., 2004; Tran et al., 2007). Indeed, Arabidopsis *CLPD* mRNA expression is induced in drought response (Kiyosue et al., 1993; Nakashima et al., 1997) and a quantitative proteomics comparison of different rice variants showed that enhanced drought tolerance was correlated to increased CLPD protein accumulation (Wu et al., 2016). Transgenic overexpression of rice CLPD in Arabidopsis plants was found to enhance drought tolerance (Mishra et al., 2016). However, the molecular role of CLPD in senescence and drought response is not yet determined.

The central question is whether CLPD indeed serves as a chaperone and delivery system for CLPPR substrates or if it functions as an unfoldase unrelated to CLP protease activity. If indeed CLPD functions as partner for the CLP protease, then CLPD should interact with the CLPPR core. Furthermore, the relationship of CLPD with CLPC1 is also not clear: do they form heterooligomers or exclusively homohexamers? Here we characterize the *in vivo* interactome of CLPD through a Walker B mutation trapping approach (Montandon et al., 2019; Rei Liao et al., 2022) and determine the phenotypic consequences of CLPD loss-of-function and overexpression in Arabidopsis. Our results show that genetic manipulation of CLPD expression impacts the rate of seedling development: i) Loss of CLPD expression results in accelerated seedling growth and development, ii) In contrast, transgenic constitutive overexpression of CLPD causes delayed development and chlorosis in seedlings and the CLPD transgene expression is suppressed in T2 and T3 generations indicative of a negative impact of CLPD overaccumulation especially in young plants. Finally, *in vivo* CLPD interactome experiments presented in this study supports that CLPD interacts (directly or indirectly) with the CLPPR protease core, as well as the CLPC chaperones. Several candidate substrates were identified in the CLPD trap, including key enzymes in riboflavin and thiamine synthesis. Integrating these new CLPD *in vivo* trapping results with previous results from CLPC1 trapping (Montandon et al., 2019; Rei Liao et al., 2022) and quantitative comparative proteomics of *clp* chaperone, adaptor and protease mutants strongly suggests that the CLP system plays a role in riboflavin and thiamin homeostasis.

## Results

### CLPD has a unique mRNA expression pattern compared to CLPC1 and CLPC2

To compare mRNA and protein accumulation of the three chloroplast CLP chaperones *CLPD*, *CLPC1* and *CLPC2*, publicly available databases were mined for expression data in *Arabidopsis*. mRNA data from ePlant (https://bar.utoronto.ca/eplant/) (Fucile et al., 2011) across development and organs shows that *CLPD* mRNA is on average about three-fold lower than *CLPC1* mRNA and is less broadly expressed than *CLPC1,* but high *CLPD* levels are found in specific organs or tissue types (Fig. 2a; Supplemental Figure S1a). While *CLPC1* mRNA levels do not vary much across leaf development, *CLPD* mRNA levels dramatically increase in senescing tissues (leaf, cauline leaf, sepals and petals), reaching ∼75% of CLPC1 levels in senescing tissue. *CLPD* mRNA is also strongly induced in osmotic stress and drought response, unlike *CLPC1* and *CLPC2* mRNA (Fig. 2a). *CLPC2* mRNA levels are, on average ∼55-fold lower than those of *CLPC1*. Interestingly, the expression pattern of *CLPC2* mRNA largely follows an inverse pattern to that of *CLPD* (Fig. 2a; Supplemental Figure S1a). The specific mRNA expression pattern of *CLPD* is unlike those of *CLPC1* and *CLPC2*, suggesting CLPD plays a unique role in the chloroplast CLP system.

**Figure 2.**
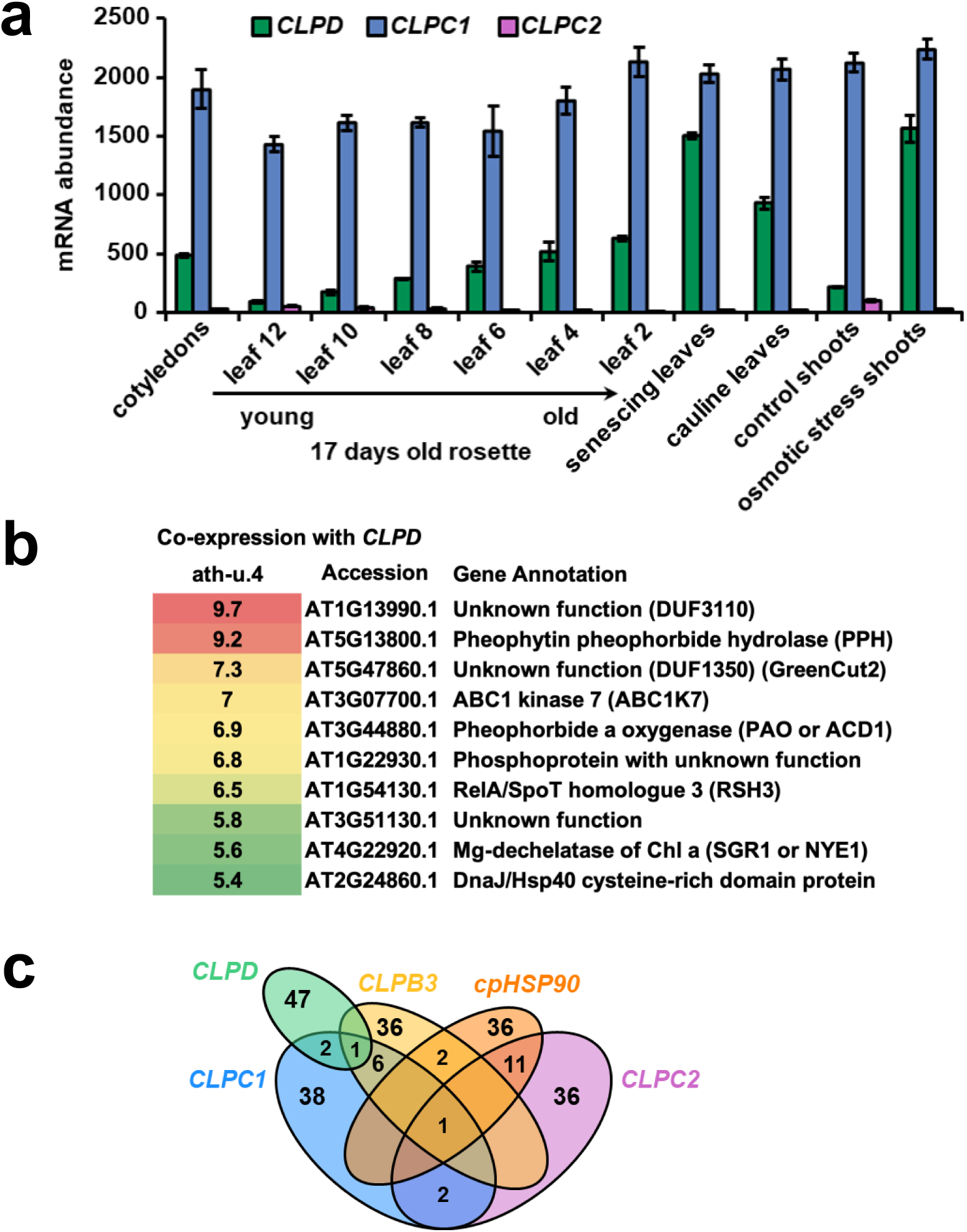
Analysis of publicly available data of mRNA expression patterns of *CLPD*, *CLPC1*, and *CLPC2*. a) mRNA expression levels from ePLANT (Fucile et al., 2011) *CLPD* (green), *CLPC1* (blue), and *CLPC2* (purple) in cotyledon and various leaf tissues as well as in shoots with and without osmotic stress (n = 3). Error bars shown are standard deviations. For cotyledon and leaf samples, plants were grown in continuous light. Leaf samples were collected from 17 days old rosettes with the exception of senescing and cauline leaves which were collected at 35 and over 21 days old, respectively (Schmid et al., 2005). b) The top 10 co-expressors of CLPD from ATTEDII. c) Venn diagram of results of mRNA-based co-expression analysis of ATTEDII u4 expression data displaying the overlaps between the top 50 co-expressors of CLPD (green), CLPC1 (blue), CLPC2 (purple), chloroplast HSP90 (cpHSP90, orange), and chloroplast CLPB3 unfoldase (yellow) (Supplemental Table S1).

mRNA co-expression data from ATTEDII (u4) (Obayashi et al., 2007) shows that all top 10 mRNA co-expressors of *CLPD* are chloroplast proteins and show highly preferential expression in senescing tissues (Fig. 2b). Three of these are involved in chlorophyll degradation, namely PHEOPHYTIN HYDROLASE (PPH;AT5G13800) (Guyer et al., 2018), PHEOPHORBIDE A OXYGENASE (PaO or ACD1; AT3G44880) (Aubry et al., 2020), and Mg dechelatase STAY-GREEN1 (SGR1 or NYE1; AT4G22920) (Sakuraba et al., 2014). The other co-expressors are (from highest to lowest co-expression coefficient): AT1G13990 (with a DUF3110, but no known functions), AT5G47860 (with a DUF1350, also named NICP1) implicated in plant immunity and ROS production (Wang et al., 2025), ABC1 kinase 7 located in thylakoid associated plastoglobules (van Wijk and Kessler, 2017), a phosphorylated protein (at S584; https://peptideatlas.org/builds/arabidopsis/) with no known functions (AT1G22930), RELA/SPOT HOMOLOG 3 (RSH3; AT1G54130) which is a ppGpp synthase implicated in defense involved in the stringent response (Masuda et al., 2008; Mizusawa et al., 2008; Maekawa et al., 2015; Honoki et al., 2018), a protein (AT2G24860) without known function, and a DnaJ domain protein (AT2G24860) also named EDS1-INTERACTING J PROTEIN1 (EIJ1) implicated in plant immunity (Liu et al., 2021).

Comparison of the top50 co-expressors of CLPC1, CLPC2, CLPD, cpHSP90 (AT2G04030) maturase and the chloroplast unfoldase CLPB3 (AT5G15450) show six unique shared co-expressors between CLPC1 and CLPB3, eleven between cpHSP90 and CLPC2, but none for CLPD with CLPC2 or cpHSP90. CLPD shares two co-expressors with just CLPC1 (PaO/ACD1 and Tic55-related protein PTC52), and one (thylakoid protease FTSH8) with both CLPC1 and CLPB3 (Fig. 2c; Supplemental Table S1). These divergent co-expression patterns strongly indicate that each of the three CLPC1,2 and CLPD chaperones have specific roles.

### CLPD mRNA and protein developmental dynamics from seedling to late flowering stage

We determined the CLPD mRNA and protein accumulation dynamics across development in rosettes under the growth conditions applied to other experiments in this study (see below). To that end, we extracted RNA and total leaf protein from wild type rosettes at specific stages of *Arabidopsis* development as defined in (Boyes et al., 2001) (Fig. 3a). qRT-PCR of *CLPD* transcripts across these developmental stages showed that *CLPD* mRNA is lowest in the initial stages of rosette development characterized by leaf production (developmental stages 1.02, 1.05, and 1.14), increases approximately five-fold in the later stages characterized by rosette expansion and emergence of the inflorescence before flowering (stages 3.70 and 5.10), and finally increases around 25-fold in flowering plants (stages 6.30 and 6.75) compared to the levels measured in the earliest stage 1.02 (Fig. 3b). We also conducted MSMS analysis of total leaf protein extracts to measure CLPD protein levels across the same developmental stages (Fig. 3c). This showed that the abundance of CLPD protein in leaves is around three-fold higher in the flowering stage (stage 6.30 and 6.75) than in plants in stage 1.14 (Fig. 3c). There was only a very modest increase in plants at stage 3.75 or 5.10 compared to stage 1.14. There was no difference in CLPD accumulation between plants stage 6.30 and 6.75, *i.e.* plants 30% of the way through flowering and plants 75% of the way through flowering. Cross-correlation of these CLPD mRNA and protein data shows an excellent linear correlation (R^2^=0.998) demonstrating that mRNA and protein abundances are highly coordinated (Fig. 3c). Our data is further supported by analysis of published quantitative mRNA and protein resources (see Supplemental Text and Supplemental Figure S1b,c).

**Figure 3.**
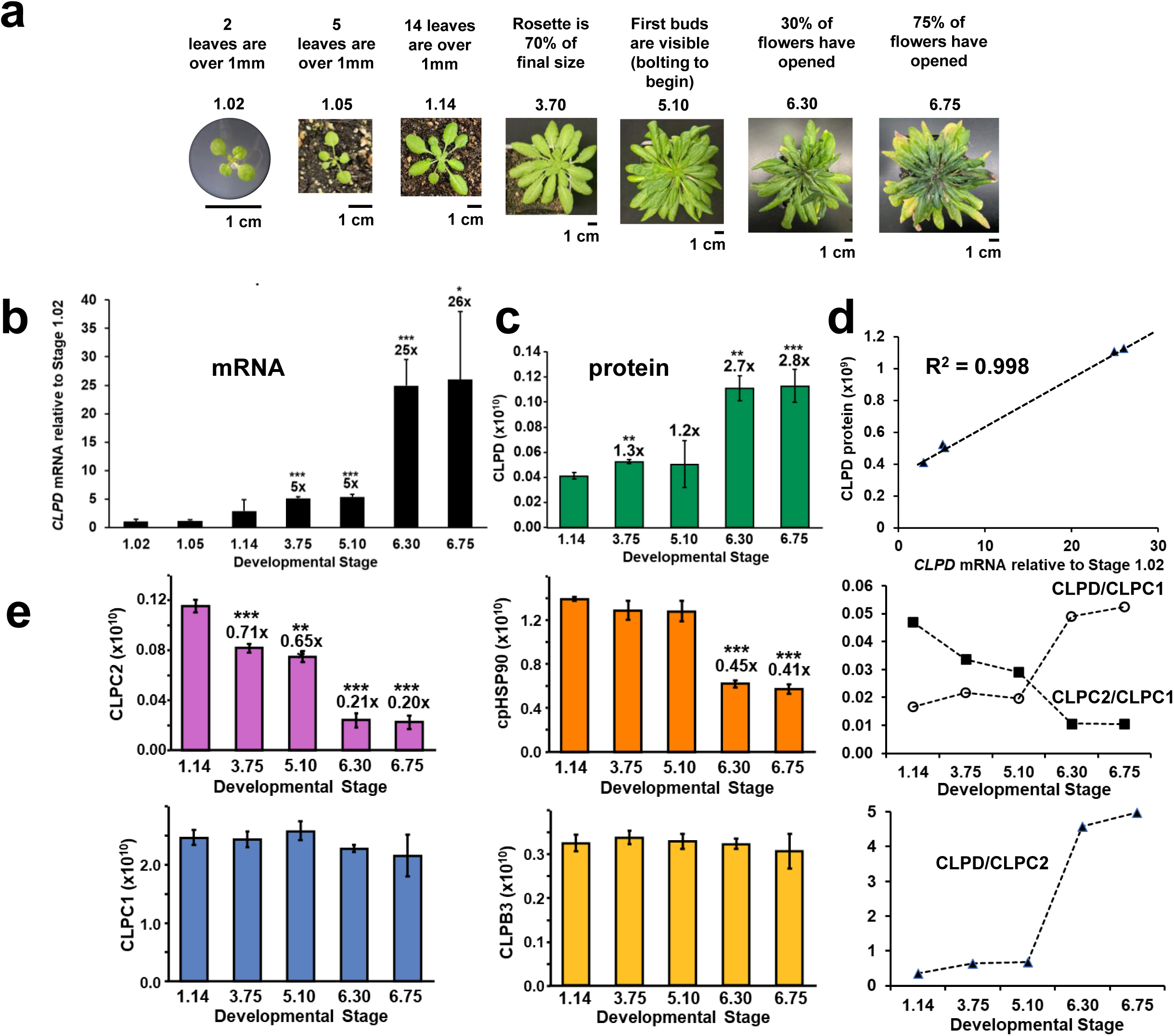
CLPD mRNA and protein levels of Arabidopsis rosettes throughout different stages of plant development. a) Developmental series of Arabidopsis rosettes with growth stages defined as in (Boyes et al., 2001). Bolts were trimmed from stage 6.30 and 6.75 plants immediately before plants were photographed. b-e) Whole leaf rosettes (with the bolts trimmed off in bolting plants) from wild type Col-0 plants were ground with liquid nitrogen and the resulting tissue sample was split for extraction of RNA and protein (three rosettes per timepoint). All p-values are calculated by two-tailed unpaired Student’s t-test and error bars shown are standard deviation, n=3. p<0.05 (*); p<0.01 (**); p<0.001 (***). b) qRT-PCR analysis of *CLPD* mRNA levels at each developmental stage is shown relative to the average level at stage 1.02 and is normalized to both the *ACTIN2* and *UBIQUITIN1* level of each sample. c-e) Protein accumulation in the same tissue measured by MSMS for c) CLPD (green) and e) CLPC2 (purple), cpHSP90 (orange), CLPC1 (blue), and CLPB3 (yellow) at each developmental stage. Equal amounts of protein of each stage (normalized to fresh weight of tissue) were separated by SDS-PAGE and the gel regions of proteins between 75-125 kDa were excised, proteins digested with trypsin and extracted peptides identified and quantified by MSMS using MS peak intensity. Protein abundances are in arbitrary units. d) Cross-correlation between relative *CLPD* mRNA levels (from panel b) and CLPD protein levels (from panel c). The right hands panels of e) display protein abundance ratios between CLPC1, CLPC2 and CLPD across development.

### Developmental dynamics of CLPD, CLPC1, CLPC2 protein abundance

Along with CLPD, we also identified and quantified (by MSMS) protein abundance levels of CLPC1 and CLPC2, as well as the major chloroplast chaperones CLPB3 and cpHSP90 (all five are within 90-100 kDa) across these five developmental stages (stages 1.14 to 6.75) (Fig. 3e). CLPC1 and CLPB3 protein levels remained quite constant through this wide developmental range, but abundance of CLPC2, and to lesser a degree cpHSP90, dropped several fold especially during the flowering stage when the rosette is undergoing natural senescence. Using these abundance values, we then calculated the abundance ratios for CLPD/CLPC1, CLPC2/CLPC1, as well as CLPD/CLPC2 (Fig. 3e). This showed that over development from seedling stage 1.14 to late flowering stage 6.75 when the rosettes are in an advanced stage of senescence, the CLPD/CLPC1 ratio 2.5 fold increased from 0.02 to 0.05. CLPC2 protein showed an opposite accumulation pattern to that of CLPD, as illustrated by the CLPC2/CLPC1 and CLPD/CLPC2 ratios. CLPC2/CLPC1 decreased ∼4-fold from 0.047 to 0.011 during development from stage 1.14 to 6.75. The CLPD/CLPC2 ratio increased 14-fold from 0.36 in the seedling stage (1.14) to 4.98 in stage 6.75 (Fig. 3e). We also note that the CLPB3/CLPC1 ratio was constant throughout development at 0.14, whereas cpHSP90/CLPC1 started out at 0.57 and decreased about 2-fold to 0.27. These data also provide approximate stoichiometries between these five chaperones, with CLPD and CLPC2 representing 1%-5% of CLPC1 levels. CLPB3 and cpHSP90 abundances are respectively ∼14% and 25%-57% of CLPC1. These general stoichiometries are very comparable to the findings of a previous analysis using large scale MSMS of isolated chloroplast stromal proteomes from mature rosettes (Zybailov et al., 2008).

### A CLPD loss-of-function mutant has accelerated growth and development

A *clpd-1* null t-DNA insertion line (Supplemental Figure S2) was previously classified (in the context of genetic interactions with CLPS1) as having no obvious visible phenotype (Nishimura et al., 2013). However, our new and careful analysis shows that seedling development is developmentally accelerated in *clpd-1* by one to two days, quantified by fresh weight of individual seedlings (Fig. 4a and b) and was not due to differences in germination rates. The accelerated developmental phenotype was still apparent after seedlings were transferred to soil and further into rosette growth, as *clpd-1* plants had more leaves than wild type plants at six weeks (Fig. 4c and d). However, there was no clear visible phenotype during senescence, no significant difference in time to emergence of the first flower buds (“bolting”) in either short day or long day conditions, and no difference in total seed weight per plant (data not shown). Hence loss of CLPD appears to remove a growth and developmental constraint during vegetative development but surprisingly does not visibly impact the natural senescence process.

**Figure 4.**
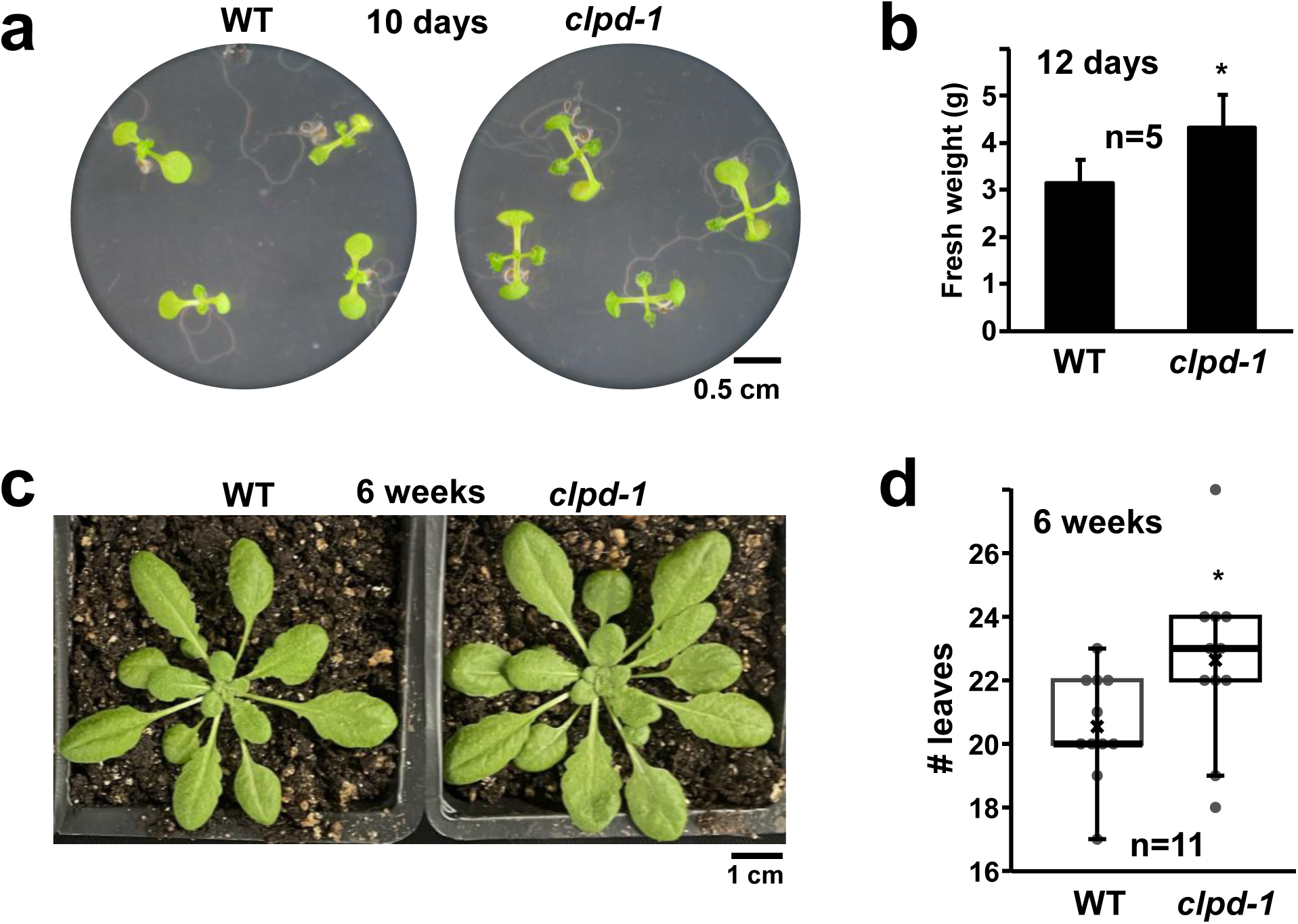
Loss of ClpD accelerated rosette development. a, b) Seedlings were grown on the same plate in short day (10 hours light) conditions. a) Light microscope images of wild type and *clpd-1* seedlings after 10 days of growth. b) Fresh weight in grams of whole wild type (WT) and *clpd-1* seedlings detached from roots measured at 12 days of growth. p<0.05 (*) determined by two-tailed unpaired Student’s t-test. Error bars are standard deviation, n=5. c,d) Plants were grown on plates in short day (10 hours light) conditions for two weeks and then transferred to growth on soil for four more weeks in long day conditions (16 hours light). c) Images of wild type and *clpd-1* plants after the total six weeks of growth. d) The number of leaves (>1 mm long) of WT and *clpd-1* plants at six weeks of growth. Individual measurements are marked with dots, n=11, median values are indicated with thicker lines, and mean values are marked with Xs. p<0.05 (*) calculated two-tailed unpaired Student’s t-tests.

### Constitutive overexpression of functional CLPD delays leaf growth and development and is silenced in T_3_ and T_4_ generations

To determine the impact of CLPD overexpression, we created two stable transgenic lines expressing wild-type CLPD with a C-terminal STREPII tag in Col-0 (WT) driven either by the genomic CLPD promoter (*gCLPD:*CLPD-STREP) or the constitutive 35S promoter (*35S*:CLPD-STREP-1,2). Transgene insertion and expression were confirmed by sequencing of cDNAs, RT- or Q-PCR as well as SDS-PAGE and immunoblotting with anti-strep serum (Supplemental figures S3 and S4). Young *gCLPD:*CLPD-STREP seedlings did not have a visible phenotype and showed no altered fresh weight or chlorophyll levels (Fig. 5a,b). The *35S*:CLPD-STREP-1,2 overexpression lines showed a delay in seedling development (∼ 1-2 days) and yellowing of newly formed leaves, quantified as decreased fresh weight and accumulation of chlorophyll a and b content (Fig. 5a,b; Supplemental Figure S3). The yellowing is only visible soon after leaf initiation and fades as the leaf grows. Delayed seedling development was not due to differences in germination time between *35S*:CLPD-STREP-1,2 and wild type plants. 35S:CLPD-STREP-1,2 T_2_ plants with detectable transgenic protein accumulation during senescence (determined by immunoblot for the STREPII tag - Supplemental Figure S4a) showed no visible senescence phenotype nor chlorophyll phenotype during flowering (data not shown). The lack of phenotypes in late development suggests that overexpressing functional CLPD is only a problem when occurring at the wrong time, *i.e.,* early in development when endogenous CLPD expression is lower. Interestingly, the homozygous T3 generation of the 35S:CLPD-STREP-1,2 lines showed loss of transgene expression both at the mRNA and protein level as the plants aged further in development (at 7 weeks old; Supplemental Figure S4b,c) likely due to silencing, but not at the seedling stage (Supplemental Figure S3c). The transgene was silenced altogether in both young and old plants in the T4 generation (data not shown).

**Figure 5.**
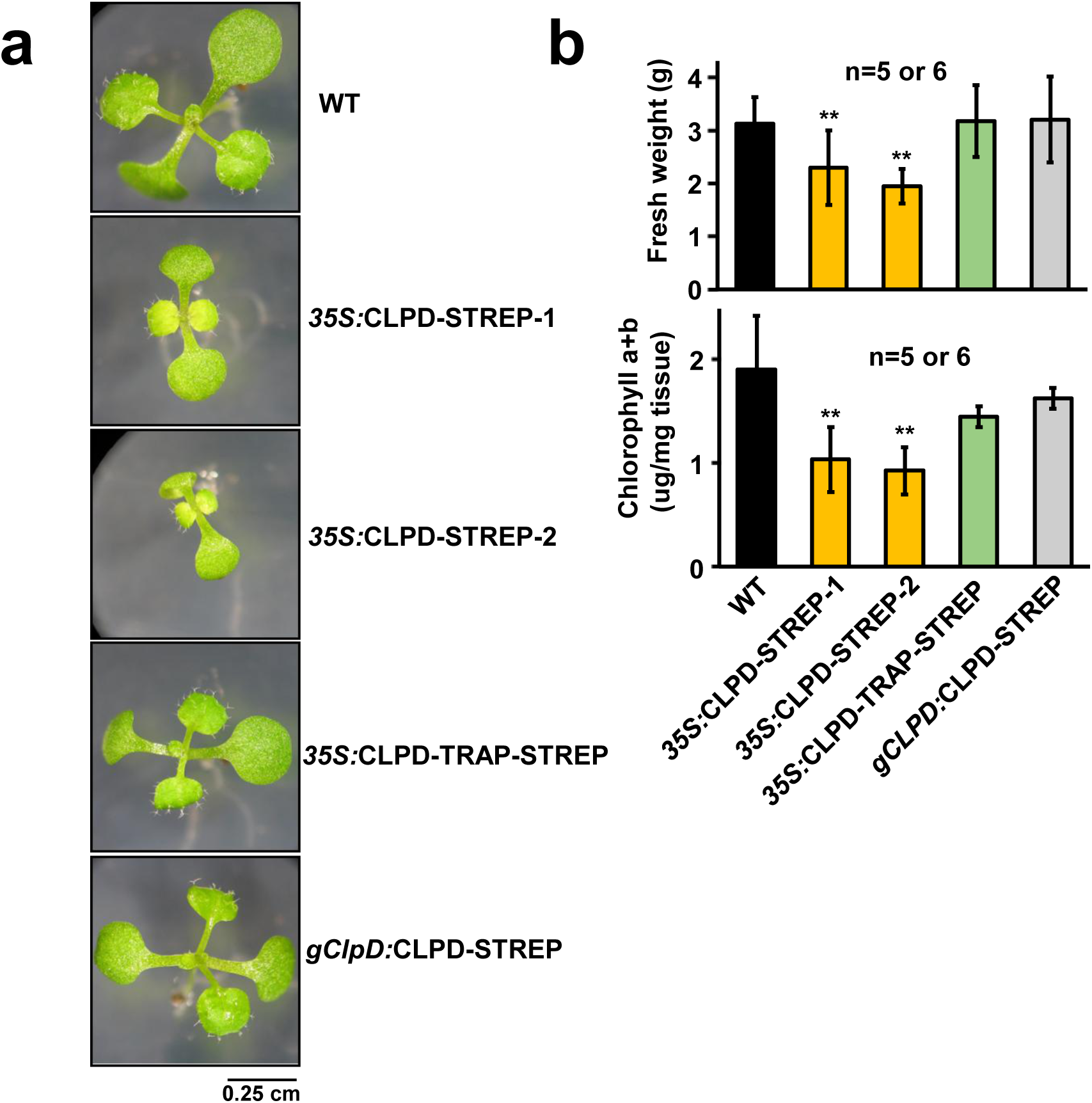
Functional CLPD overexpression, but not CLPD-TRAP overexpression, leads to delayed development and decreased chlorophyll levels. Wild type seedlings were compared to seedlings from homozygous transgenic lines overexpressing either functional CLPD (CLPD-STREP-1,2 with 1 and 2 representing independent transformants) or Walker B mutated CLPD (CLPD-TRAP-STREP) under the 35S or genomic *CLPD* promoter (*gCLPD*) in the wild type background. Seedlings were grown on the same plate in short day (10 hours light) conditions. a) Light microscope images of representative seedlings from each transgenic line and wild type grown for 12 days. b) Fresh weight and chlorophyll a and b content (normalized to fresh weight) of wild type, *35S:*CLPD-STREP-1, *35S:*CLPD-STREP-2, *35S:*CLPD-TRAP-STREP, and *gCLPD:*CLPD-STREP seedlings were measured after twelve days of growth. Error bars are standard deviation. P values compare the different CLPD overexpression lines to wild type and are calculated by two-tailed Student’s t-test. p<0.01 (**) and p<0.001 (***).

### Generation of CLPD-TRAP lines to identify CLPD protein interactors

To identify CLPD interactors, determine if CLPD might form complexes with the CLP protease core complex, and possibly capture CLPD substrates, we employed a similar ‘trapping’ strategy as we previously successfully used for CLPC1 with 35S-driven expression of CLPC1 transgenes (Montandon et al., 2019; Rei Liao et al., 2022). This strategy makes use of the conserved glutamate residues in the Walker B motif of each AAA domain of the chaperone that are critical for ATP hydrolysis. Mutation of these residues (E388A and E737A in CLPD) prevents the chaperone from carrying out ATP-dependent substrate unfolding but does not affect substrate binding, thus allowing for affinity purification and MSMS analysis without release and degradation of interactors. We generated a stable transgenic line in the wild type Col-0 background transformed with a construct coding for CLPD-TRAP (E388A and E737A) with a C-terminal STREPII tag under the constitutive 35S promoter (*35S*:CLPD-TRAP-STREP). The CLPD-TRAP-STREP protein was easily detectable by SDS-PAGE and immunoblotting (Supplemental Figure S5a). While *35S*:CLPD-TRAP-STREP has no seedling phenotype (Fig. 5a,b), the plants grown under short day conditions (10 h light/14 h dark) develop a patchy yellowing and downward curling leaves phenotype beyond stage 3.50 (characterized by rosette expansion) that worsens as the plants age (Supplemental Figure S5b). However, when the *35S*:CLPD-TRAP-STREP line plants were grown under long-day conditions (16 h light/8 h dark), these late-stage phenotypes did not emerge (Fig. 6a). Hence, the phenotypic consequences of constitutive overexpression of CLPD-TRAP are impacted by the lengths of the day/light periods. We also successfully generated a *gCLPD*:CLPD-TRAP-STREP transgenic line (with the CLPD genomic promoter). CLPD-TRAP-STREP protein level in this genomic promoter line was relatively low but detectable by immunoblotting, and no visible phenotype was observed (Supplemental Fig S6).

**Figure 6.**
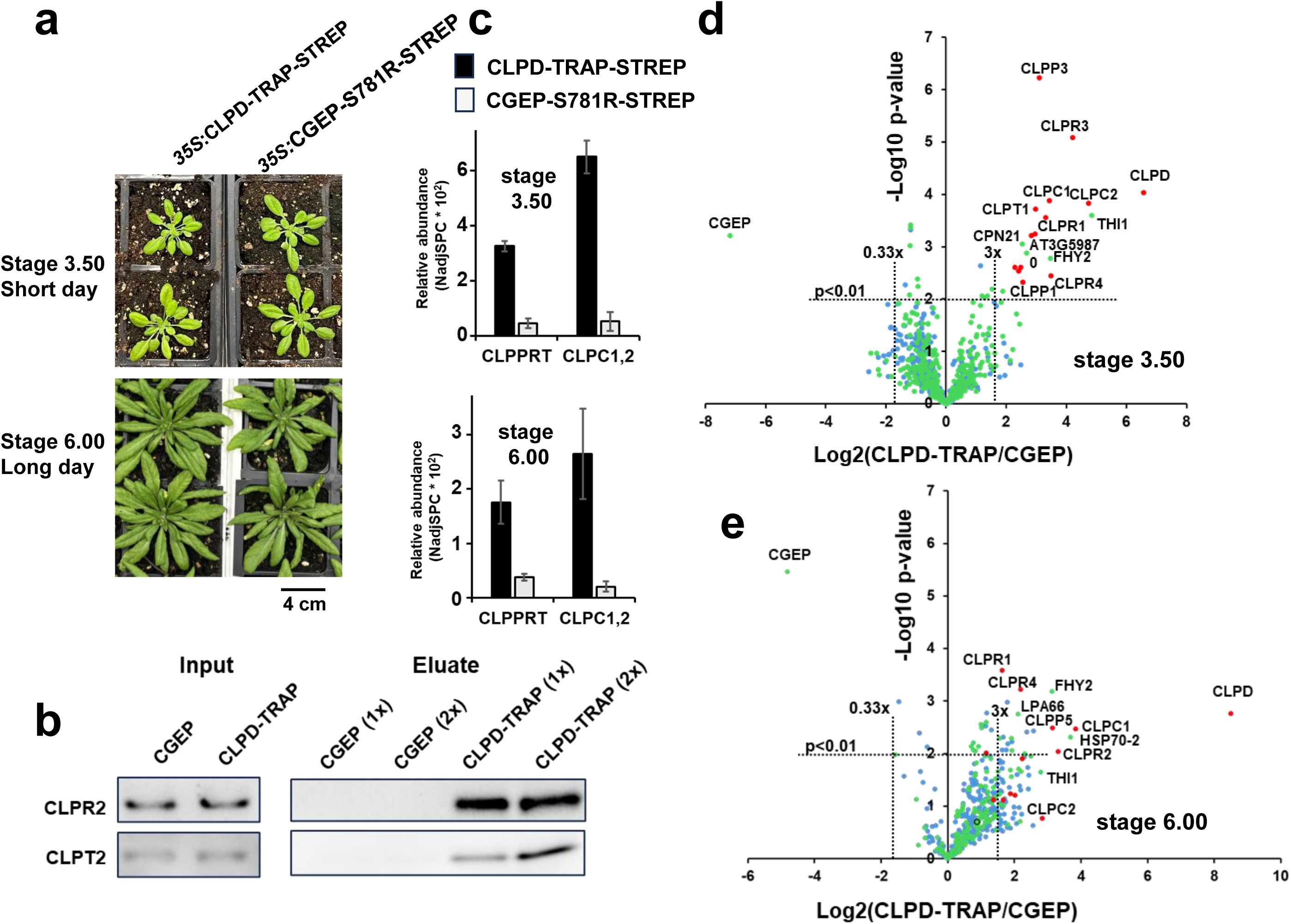
*In vivo* trapping experiments with *35S:*CLPD-TRAP and *35S*:CGEP-S781R lines. Point mutations in each Walker B domain (E388A and E737A) prevent CLPD driven ATP hydrolysis activity resulting in a block of substrate unfolding and subsequent degradation while stabilizing CLPD interactors. The homozygous *35S*:CGEP-S781R-STREP line (Bhuiyan et al., 2020) was used as a negative control. a) *35S:*CLPD-TRAP and *35S:*CGEP-S781R-STREP plants used for trapping at stage 3.50 grown in 10 hours light period (top) or at stage 6.00 grown in 16 hours light period (bottom). Plants were photographed immediately prior to rosette tissue collection for trapping experiments. b) Immunoblot of inputs (*35S:*cGEP-S781R-STREP and *35S:*CLPD-TRAP-STREP soluble protein extracts, left) and eluates (loaded in a titration of 1x or 2x volume loaded, right) from the “before bolting” trapping experiment using anti-CLPR2 or anti-CLPT2 sera. c) Comparison of average normalized intensity for CLPC1,2 chaperone proteins as well as CLPPR core proteins and CLPT1,2 accessory proteins in MSMS analysis of *35S:*CLPD-TRAP-STREP and *35S:*CGEP-S781R-STREP eluates for the before bolting (top) and after bolting (bottom) trapping experiments. d) Volcano plots of proteins identified by MSMS in the *35S:*CLPD-TRAP-STREP and *35S:*cGEP-S781R-STREP eluates affinity purified from Arabidopsis before bolting in stage 3.50 (top) and after bolting in stage 6.00 (bottom). Red points are CLP core, CLP chaperone, adaptor, or CLP accessory proteins. Green points are proteins with confirmed plastid localization (as annotated in PPDB). Blue points are proteins with non-plastid or unknown localization. Horizontal dashed lines indicate p values of 0.01 and vertical dashed lines indicate thresholds of 3-fold enrichment either in the *35S:*CLPD-TRAP-STREP or *35S:*CGEP-S781R-STREP eluates.

Attempts to purify transgenic CLPD protein from *gCLPD:*CLPD-STREP and *gCLPD:*CLPD-TRAP-STREP lines by streptactin affinity purification resulted in insufficient yields for MSMS-based identification of interactors (data not shown). We therefore focused on the *35S*:CLPD-TRAP-STREP plants for the CLPD interactome analysis as transgenic CLPD-TRAP-STREP levels were sufficient for affinity enrichment and MSMS-based identification. Trapping experiments were performed on plants grown in conditions designed to minimize impact of development phenotypes. A trapping experiment was conducted using plants grown in a short-day light period with tissue harvested before bolting at growth stage 3.50 before the short-day light period-dependent *35S*:CLPD-TRAP-STREP phenotype becomes highly pronounced (Fig. 6a). A second trapping experiment was conducted using plants grown in long-day conditions to prevent the short day phenotype, and tissue was collected soon after bolting at growth stage 6.00 (Fig. 6a). As negative control for CLPD protein interactors, we used our previously published transgenic line that expresses a catalytically inactive variant (S781R) of CHLOROPLAST GLUTAMYL ENDOPEPTIDASE (CGEP), a stromal chloroplast peptidase functionally unrelated to the CLP system (Bhuiyan et al., 2020), with a C-terminal StrepII tag under the *35S* promoter (*35S*:CGEP-S781R-STREP). This *35S*:CGEP-S781R-STREP line has no visible phenotype (Fig. 6a) and CGEP-S781R does not have known stable protein interactors (Bhuiyan et al., 2020).

### *In vivo* CLPD trapping demonstrates association with the CLPPR core complex and identifies additional protein interactors

MSMS analysis of the affinity eluates from *35S*:CLPD-TRAP and *35S*:CGEP-S781R plants grown under short-day conditions and harvested before bolting (stage 3.50) identified 751 proteins after grouping of proteins with a high percentage of shared matched spectra and application of a minimum matched spectral threshold (>6 adjSPC) (Supplemental Dataset S1). These 751 proteins included all 15 chloroplast CLP proteins including CLPF, but except CLPS1 because CLPS1 is hard to identify by MS due to its small size and low number of suitable tryptic peptides (Nishimura et al., 2013). The CLPPRT core, CLPC1,2 and CLPD were respectively 7-, 12- and 95-fold enriched in the CLPD affinity eluates compared to CGEP eluates (Fig. 6c – upper panel). Conversely, CGEP was 145-fold enriched in the CGEP eluates (Table 1). This shows that CLPD interacts with both the CLP protease core and CLPC1,2. The volcano plot for CLPD-TRAP/CGEP ratios shows the distribution of the two baits (CGEP and CLPD), the CLP proteins, and other plastid and non-plastid proteins (Fig. 6d) (Supplemental Dataset 2). At p<0.01 and at least three-fold change this identified CGEP and CLPD as well as 17 other proteins all of which were plastid localized (Table 1). This included all CLP proteins except CLPF and CLPS1, as well as chaperonin 20 (CPN20), thiazole biosynthetic enzyme 1 (THI1; AT5G54770) (Chabregas et al., 2003), ARPP phosphatase (FHY2/PYRP2; AT4G11570) involved in riboflavin biosynthesis (Sa et al., 2016), and two proteins with unknown functions AT3G59870 and AT5G45170. AT3G59870 has one homolog AT2G43945 which is also plastid-localized but neither function has been determined and functional domain predictions are inconclusive. AT5G45170 belongs to the haloacid dehalogenase family, similar to FHY2/PYRP2, but unlike FHY2/PYRP2 it has no substrate affinity for ARPP and the function of AT5G45170 remains unknown (Sa et al., 2016). CPN20 has been identified as an interactor to the CLP protease core in several publications for Arabidopsis (Kim et al., 2015; Rei Liao et al., 2022; van Wijk, 2024) as well as Chlamydomonas (Ning Wang et al., 2021). We do note while CLPF did not meet the statistical criterium for CLPD-TRAP enrichment, CLPF was identified in each CLPD-TRAP replicate with a total of 15 matched spectra and identified with only 2 spectra in the CGEP eluates; this suggests that also CLPF is part of the CLPD interactome.

**Table 1.**
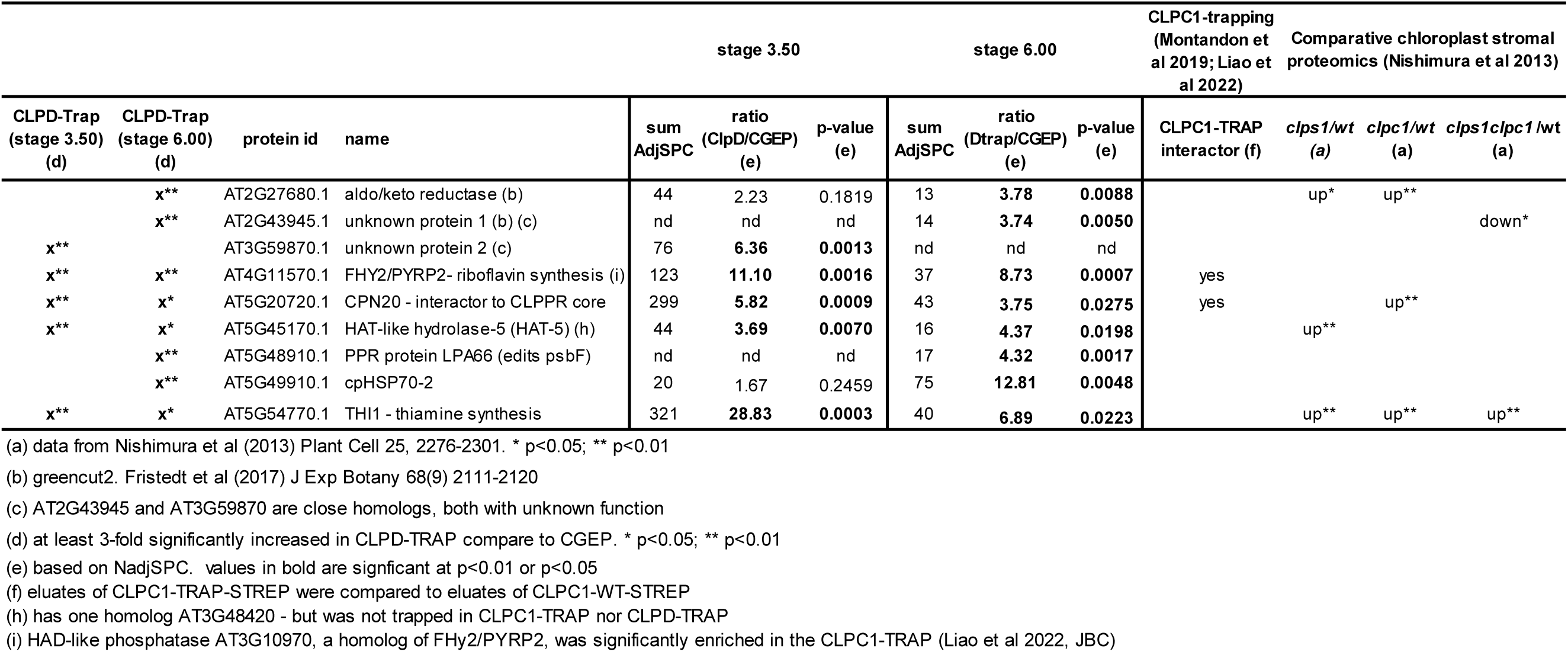
Proteins identified in CLPD-TRAP at stage 3.50 and/or stage 6.00 (p<0.01)

In the post-bolting experiment (stage 6.00), MSMS analysis of the affinity eluates of *35S*:CLPD-TRAP and *35S*:CGEP-S781R identified 488 proteins and protein groups (Supplemental Dataset S1). This included 14 chloroplast CLP proteins with only CLPP3 and CLPS1 missing. The CLPPRT core, CLPC1,2 and CLPD were respectively 5-, 13- and 361-fold enriched in the CLPD affinity eluates compared to CGEP eluates (Fig. 6c – lower panel) (based on NadjSPC). Conversely, CGEP was 28-fold enriched in the CGEP eluates. This again shows that CLPD interacts with both the CLP protease core and CLPC1,2. A volcano plot for CLPD-TRAP/CGEP ratios shows the distribution of CGEP, CLPD, the CLP proteins as well as other plastid and non-plastid proteins (Fig. 6e). 19 proteins were identified with p<0.01 and at least 3-fold enrichment, 11 of which were plastid localized, including 6 CLP proteins (CLPD, CLPC1, CLPR3, CLPR4, CLPP5) and CGEP (Table 1). The non-CLP chloroplast proteins were: cpHSP70-2 (AT5G49910), FHY2/PYRP2, PPR protein LPA66 (AT5G48910) which helps editing of chloroplast-encoded psbF mRNA (Cai et al., 2009), as well as AT2G27680 (aldo/keto reductase) and AT2G43945 with unknown functions.

As shown in Table 1, the two trapping experiments identified in total nine chloroplast proteins (in addition to most CLP proteins) as candidate interactors (at our relative stringent criteria) of CLPD, with FHY2/PYRP2 identified in both at p<0.01 and HAT-5 and THI1 in both at either p<0.01 or p<0.05. Interestingly, AT2G43945 (unknown protein 1) which was enriched in stage 6.00 has a single close homolog AT3G59870 (unknown protein 2), which was enriched in stage 3.50 (Table 1); Verification of the matched peptides showed that these are specific identifications for each experiment.

## Discussion

### The importance of developmentally regulating CLPD protein abundance

Close examination of mRNA and protein data from different sources and new experimental data from this study show that CLPD protein levels positively correlate with CLPD mRNA levels, both increasing in senescing tissues. This conclusively resolves the earlier dispute about the correlation between CLPD mRNA and protein levels (Weaver et al., 1999). Complete loss of CLPD expression (the *clpd-1* null line) accelerated seedling development, but the underlying mechanism is not clear. However, even in the cotyledons and young leaves, CLPD protein is already present and its role in development-associated processes is likely not constrained by the extent of cellular senescence. It is well appreciated that senescence is more than simply a degradation and remobilization process, and involves cellular responses at transcriptional, translational, metabolic and hormonal levels (Kim et al., 2018; Zhai et al., 2026). Surprisingly, loss of CLPD (in *clpd-1*) did not result in quantifiable senescence phenotypes, even if the CLPD protein level in WT increases several fold during senescence. In contrast, constitutive over-expression of functional CLPD caused delayed growth and chlorosis in the seedling stage indicative of partial inhibition of chloroplast biogenesis during early stages of leaf development by excess CLPD. This CLPD overexpression seedling phenotype is dependent on CLPD ATPase/unfoldase activity, as it did not occur in seedlings expressing CLPD-TRAP (unable to unfold substrates) under the same constitutive promoter. The constitutively expressed functional CLPD transgene was silenced in later generations of transgenic lines. Together, this shows that CLPD protein levels must be carefully controlled as a function of leaf and rosette development possibly to avoid mis-timed degradation of CLPD-specific substrates.

These results for CLPD are very different from those observed for CLPC1, since overexpressing CLPC1 in Arabidopsis has no phenotype (Montandon et al., 2019), whereas loss of CLPC1 results in a strong phenotype of visibly pale, small plants with delayed growth and development and proteostasis stress (Nakabayashi et al., 1999; Nishimura et al., 2013; Sjogren et al., 2014). CLPC2 has a very high amino acid sequence identity to CLPC1 and has mostly a redundant function to CLPC1 as supported by the observations that *clpc1clpc2* double mutants are embryo lethal, and that overexpression of CLPC2 in *clpc1* null suppresses most of the pronounced *clpc1* virescent phenotype (Kovacheva et al., 2007). However, in addition to the CLPC1-type function, CLPC2 also appears to have a specific function in biotic stress defense (Lopez et al., 2024). Additionally, we showed in this study that the CLPC2 and CLPD mRNA and protein accumulation patterns are inversely correlated during development (see also (Sjogren et al., 2014)), suggesting that CLPC2 and CLPD each have specialized and perhaps complementary or antagonistic roles, most likely also requiring CLPC1 (see below). Together, these phenotypic findings indicate functional and regulatory specialization of the CLPC1,2 and CLPD chaperones.

### CLPD co-purifies with the CLP protease core

It has not been formally established that CLPD functions as substrate delivery chaperone and interactor to the CLPPR protease core complex. CLPD has only ∼50% amino acid sequence identity compared to CLPC1,2, but it does contain the critical IGF motif (the P loop) that is essential for docking onto the adaxial side of CLP complexes in all studied bacterial CLPC-CLP protease systems (Azinas and Carroni, 2026). However, unlike CLPC1,2, CLPD lacks a conserved R motif (Fig. 1) implicated in binding to the CLP protease core in cyanobacteria but not other bacterial species (Tryggvesson et al., 2012). Removing the conserved R motif from cyanobacterial *Synechococcus elongatus* CLPC prevents *in vitro* substrate delivery to the CLPP3R core and allows binding to a homomeric bacterial CLPP core (Tryggvesson et al., 2012). The significance of the R motif of CLPC1,2 in Arabidopsis, and angiosperms in general, is not known. Using an *in vivo* trapping approach in which a modified CLPD lacking ATP hydrolysis capacity was stably overexpressed, followed by affinity purification of the CLPD protein and MSMS analysis of its interactors, this study shows that CLPD co-purified with all proteins of the CLPPRT protease. Co-purification does not formally prove the functional docking on CLPD onto the adaxial side of the CLP protease via the well-established (both cryo-EM and X-ray crystallography resolved structures) and conserved structural interaction between hexameric CLP chaperones and tetradecameric CLP protease core complexes. It is thus possible, but in our view unlikely, that the CLPD-CLP protease interactions are indirect, perhaps via CLPC1. Because CLPD is of low abundance and the total yield of CLPD-TRAP complexes from transgenic Arabidopsis results only in sub-microgram amounts, it is not feasible to determine the oligomeric organization of CLPD-TRAP complexes. Extensive efforts to obtain such structural information with the far more abundant CLPC1-TRAP complexes from transgenic CLPC1-TRAP lines have not been successful (Annis, Ravenburg, Ehrlich and van Wijk, unpublished), illustrating the challenge for CLPD-TRAP. Hence, whereas this study has not formally proven that CLPD directly interact with the CLP protease core, the collective information for CLP chaperone-protease systems makes this the most plausible and simplest explanation. In vitro reconstitution with recombinant CLPD and reconstituted protease complexes from recombinant P, R and T subunits will be needed to answer this question more definitively.

### Current understanding of oligomerization of Arabidopsis CLPC1, CLPC2 and CLPD from in vitro and in vivo experimentation

Neither this study nor prior publications provide direct experimental data for the *in vivo* hetero- or homo-oligomerization state of CLPD. *In vitro* analysis with purified recombinant CLPD or CLPC2 showed that each can form homodimers and that upon addition of ATP homo hexamers were formed (Rosano et al., 2011). Size exclusion chromatography and small-angle X-ray scattering (SAXS) with highly purified recombinant CLPC1 showed that CLPC1 forms hexamers in the presence of 2 mM ATP and MgCl_2_, is monomeric without ATP, and no other oligomeric states were observed (Jagdev et al., 2021). Together this shows that the chloroplast CLPC/D chaperone can self-assemble into homo-hexamers without the need of any adaptor, similar to CLPC homologs in *Mycobacterium tuberculosis* and *Synechococcus elongatus* (see review (Annis et al., 2024)). This is different from the CLPC homologs in *Bacillus subtilus* and *Staphylococcus aureus* which do require adaptors such as MecA to form hexamers; without such adaptors they can form inactive dodecamers and even larger oligomeric states mediated through their Uvr motifs in the middle domain (reviewed in (Annis et al., 2024; Azinas and Carroni, 2026). There are no published reports in which recombinant CLPC1, CLPC2 and/or CLPD were mixed with each other to determine possible hetero oligomeric states. Previous analyses of isolated chloroplast stroma with 2D native gels and size exclusion chromatography are consistent with the model that the Arabidopsis chloroplast CLP chaperones form both dimers and hexamers in vivo (Peltier et al., 2006) (Olinares et al., 2010). A third paper also used size-exclusion chromatography (10 mM NaCl and 10 mM MgCl_2_ at pH 8.0) without ATP followed by immunoblotting for CLPC1,2 for chloroplast stromal proteome from wt and *clpf* (Nishimura et al., 2015). CLPC1,2 were each observed between 100-600 kDa in wt but in a narrower mass range from ∼100 - ∼200 kDa in *clpf*. These combined *in vitro* and *in vivo* studies indicate that the CLPC1,2 and CLPD chaperone form oligomers, as expected, but it is not known if heteromers are formed.

### Relative abundances of CLPC1,2 and CLPD in the denatured proteome and in the interactomes of CLPC1-TRAP and CLPD-TRAP

The abundance ratios of CLPD, CLPC1 and CLPC2 determined in this current study by MSMS of denatured leaf protein extracts across different developmental stages show that CLPD represents between 1.5% and 5% of total CLPC1,2 abundance, in general agreement with a large scale MSMS of denatured stromal proteomes of Arabidopsis full grown rosettes (Zybailov et al., 2008). Additionally, our MSMS data show that CLPD and CLPC2 have a comparable abundance range across development (from seedling to senescence) but with completely opposing patterns of abundance throughout development. This is also supported by an independent analysis using immunoblotting as detection tool (Sjogren et al., 2014). In other words, there are on average between ∼2-5 molecules of CLPD for every ∼100 CLPC1 molecules present in the chloroplast proteome. CLPC1 and CLPC2 were among the most abundant and enriched interactors in our CLPD trapping experiments. Conversely, CLPD and CLPC2 were the most abundant interactors of CLPC1 in our previously published CLPC1 trapping experiment, despite their relatively low abundance in the chloroplast proteome (Rei Liao et al., 2022). These general abundance ratios in the denatured proteome (low for CLPC2 and CLPD) and the CLPC1 and CLPD-trapping experiments (high for CLPC2 and CLPD) strongly suggest that CLPD interacts with CLPC1 and CLPC2, in addition to interacting with the CLP protease core complex, as discussed above.

### Working models for CLPD oligomerization and functions

Based on our knowledge of homologous CLP chaperone structures in bacteria and structural predictions using AlphaFold (Abramson et al., 2024), there is no reason to believe that the chloroplast CLP chaperones cannot form hetero-oligomeric hexamers. CLP chaperone hexamers are largely held together by interactions between the AAA+ domains (Lee et al., 2003; Carroni et al., 2017; Rizo et al., 2019; Xu et al., 2024), which are highly conserved in CLPC1,2 and CLPD Indeed, the collective data presented here is most fitting with mixed CLPC/D chaperone rings. Incorporating CLPD into CLPC rings could help offset the seemingly different kinetic and regulatory properties of CLPD. CLPD has a lower *in vitro* ATPase rate (Rosano et al., 2011) and lacks the UVR motif that is used for chaperone activation in bacteria (Carroni et al., 2017; Annis et al., 2024). CLPD also has an N-domain but it has a different structure than the CLPC N-domain which likely impacts the affinity and recruitment of substrates and even adaptors (Mohapatra et al., 2017). By our measurement, CLPC1 protein levels are around 20 times higher than CLPD protein levels in senescent plants. By this estimate, about 30% of CLPC1 hexamers could contain a single CLPD subunit during senescence if oligomerization is determined purely by stoichiometry; this number would be reduced to ∼10% in younger plants. Future experiments should be conducted to test the working model of chloroplast chaperone heterooligomers and might require high resolution structural analysis by cryo-EM and *in vitro* reconstitution and activity essays.

### CLPD interactors, substrates and role of the CLP system in thiamin and riboflavin metabolism

The CLPD trapping experiments identified nine enriched CLPD interacting proteins, in addition to most of the CLP proteins. As illustrated in Table 1, four of these (aldo/keto reductase, HAD-like hydrolase-5, CPN20, THI1) were statistically significantly upregulated in *clps1*, *clpc1 and/or clps1clpc1* (Nishimura et al., 2013) whereas THI1 was also upregulated in the *clpr2-1* mutant (Zybailov et al., 2009).

The ARPP phosphatase PYRP2 (also named FHY2) (a HAD-like hydrolase) required for riboflavin (vitamin b1) synthesis (Sa et al., 2016) (Fig. 7a) was identified in the CLPD-traps in both stage 3.50 and 6.00, and it was also highly enriched in our previous CLPC1 trapping studies (Montandon et al., 2019; Rei Liao et al., 2022). Riboflavin is the precursor for the essential cofactors FAD and FMN, and riboflavin also induces priming of defense responses (Eggers et al., 2021; Tsitsekian et al., 2025) (Fig. 7a). Degradation of PYRP2 in the riboflavin pathway by the CLP system would allow down-regulation of riboflavin production and thus provide a mechanism for control (Fig. 7a). In our previous CLPC1 trapping study we also identified a homolog of PYRP2 (AT3G10970) but it has no ARP phosphatase activity and its substrate is not known (Sa et al., 2016). Furthermore, PYRP2 as well as RIBA1 (AT5G64300) responsible for key steps in the riboflavin pathway (Fig. 7a), are both many fold increased in *clpc1* (Nishimura et al., 2013) further supporting regulatory control of this pathway by the CLP system.

**Figure 7.**
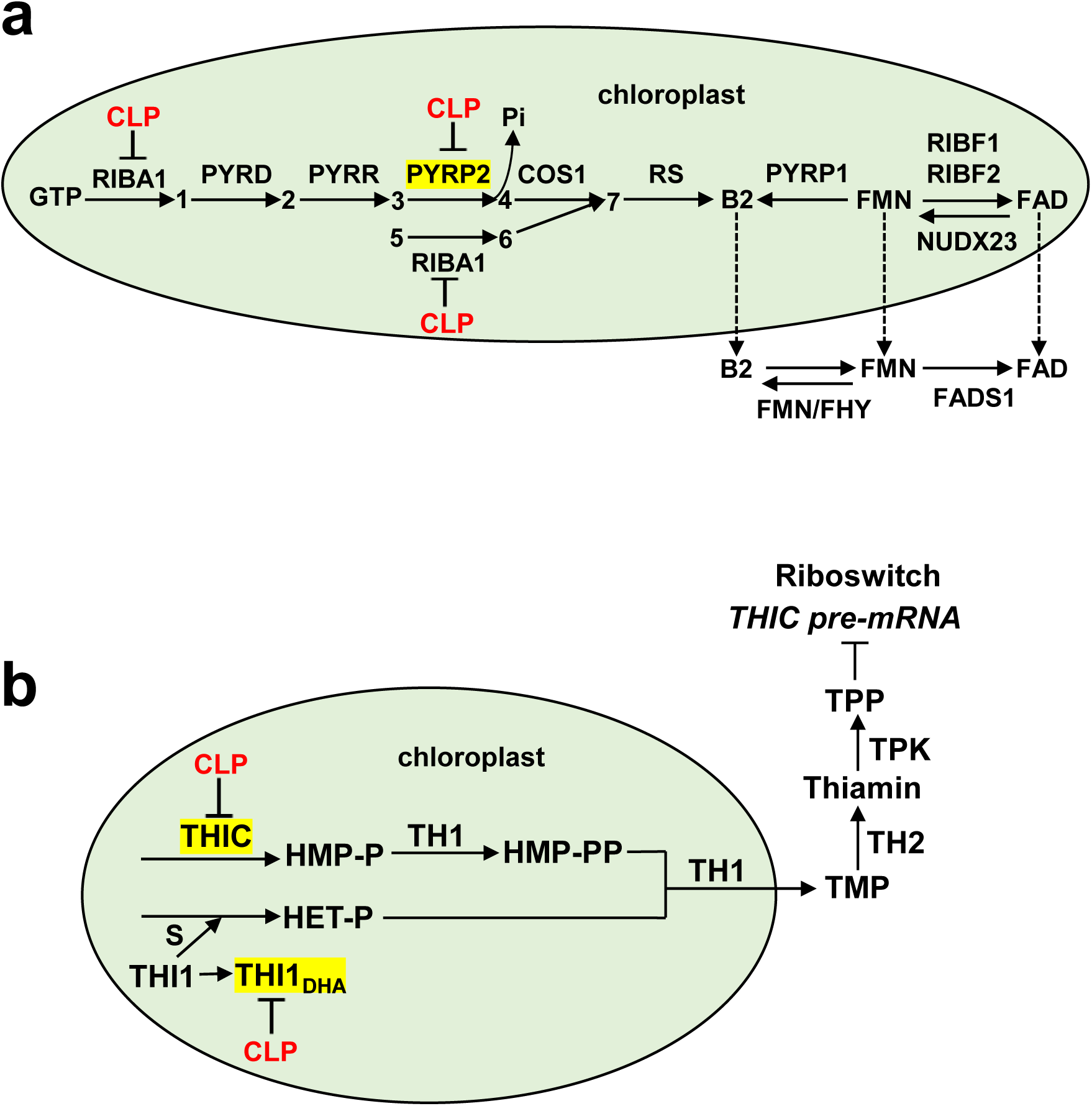
CLP involvement of regulation specific steps in the riboflavin (a) and thiamin (b) metabolic pathways. Proteins marked up in yellow are highly enriched in the CLPD trap (PYRP2 and THI1) and/or CLPC1 trap (PYRP2 and THIC), whereas RIBA1, PYRP2, THIC and THI1 are all highly increased in the *clpc1* null mutant indicative of degradation by the CLP system. Pathway and their enzymes are based on (Tsitsekian et al., 2025) and (Strobbe et al., 2021; Fitzpatrick, 2024) for riboflavin and thiamin, respectively. (1) 2,5-Diamino-6-ribosylamino-4(3H)-pyrimidinone 5’-phosphate, (2) 5-Amino-6-ribosylamino-2,4(1H,3H)-pyrimidinedione 5’-phosphate, (3) 5-Amino-6-ribitylamino-2,4(1H,3H)-pyrimidinedione 5’-phosphate (ARPP), (4) 5-Amino-6-ribitylamino-2,4(1H,3H)-pyrimidinedione, (5) D-Ribulose-5-phosphate,(6) 3,4Dihydroxybutanone 4-phosphate, (7) 6,7-Dimethyl-8-(D-ribityl)lumanize. For Arabidopsis gene identifiers and full names see (Tsitsekian et al., 2025) and (Strobbe et al., 2021).

As depicted in figure 7b, THI1 (AT55G54770), THIC (AT2G29630) and TH1 (AT1G22940) together produce the direct thiamin precursor thiamin monophosphate (TMP) in the chloroplast, which is then exported to the cytosol to produce thiamin pyrophosphate (TPP) serving as cofactor in multiple important enzymes involved in glycolysis, the TCA cycle, nucleotide metabolism and synthesis of branched chain amino acids (Dong et al., 2015; Strobbe et al., 2021; Fitzpatrick, 2024; Wang et al., 2025). The relative abundance of THI1:THIC:TH1 is about 15:3:1 based on the number of matched MSMS spectra in the Arabidopsis PeptideAtlas (https://peptideatlas.org/builds/arabidopsis/).

THI1 was identified as a CLPD interactor at stage 3.50 (∼29 fold enriched in the CLPD trap) and at stage 6.00 (∼7 fold enriched in the CLPD trap) (Table 1). THI1 was not identified as an interactor of the CLPC1-TRAP (which was compared to enrichment in CLPC1-WT eluates) in previous studies (Montandon et al., 2019; Rei Liao et al., 2022), but THI1 levels increased (p<0.01) in the chloroplast stromal proteome of *clps1, clpc1* and *clps1clpc1* mutants, by 2.3, 17 and 4.3 fold respectively (Nishimura et al., 2013). THI1 was also upregulated in the *clpr2-1* mutant (4-7 fold) in seedlings (stage 1.07 and 1.14) (Zybailov et al., 2009). THI1 has been reported to have an estimated half-life of about five hours (Li et al., 2017) which correlates with its function as a ‘suicide enzyme’ that is irrevocably damaged after a single turnover (Joshi et al., 2021; Fitzpatrick, 2024). During the catalytic reaction, THI1 donates a sulfur atom from one of its backbone cysteine residues which irreversibly converts the cysteine into dehydroalanine (DHA) inactivating its catalytic ability (Chatterjee et al., 2011; Joshi et al., 2021). Since there is no known repair mechanism, THI1_DHA_ is permanently inactivated and must be degraded, consistent with the short half-life of THI1. Indeed, most of the THI1 in Arabidopsis leaves accumulates in the form of THI1_DHA_ as determined by MSMS analysis (Joshi et al., 2021). It is unclear how THI1 is recognized as substrate for degradation by the CLP system, but formation of DHA is believed to form crosslinks with proteins, likely marking it for degradation (Paredes et al., 2023). THIC was found to be highly enriched in the CLPC1-TRAP (Montandon et al., 2019; Rei Liao et al., 2022) and THIC levels increased in the *clpc1* and *clps1clpc1* mutants by respectively 7 and 2.4 fold (Nishimura et al., 2013). We did observe THIC in the CLPD-TRAP, but because the number of matched spectra was low in the CLPD trap (but detected in each replicate), this did not pass our statistical significance threshold (even if it was not detected in the negative control CGEP). Both THIC and THI1 were also increased in *clp* protease RNAi lines in tobacco consistent with our observations in Arabidopsis (Moreno et al., 2018; Rodriguez-Concepcion et al., 2019). Our current CLPD and prior CLPC1 trapping studies as well as comparative proteomics of *clp* mutants, collectively suggest that THIC and THI1 are substrates for the CLP protease system (Fig. 7b).

CPN20 was enriched in our CLPD trapping data (Table 1) as well as our previous CLPC1 trapping experiments (Montandon et al., 2019; Rei Liao et al., 2022). CPN20 was also highly enriched in CLP protease complexes purified via transgenic tagged CLPP5 (Liao et al., 2018) and also of transgenic tagged CLPT1,2 (Kim et al., 2015). A cryo-EM structure of the green algae *Chlamydomonas reinhardtii* CLPPR core revealed that CPN20 co-purified with the core and docked to the P ring but the role of this tight interaction between CPN20 and the CLP protease is not clear (Wang et al., 2021). Our findings add to the growing evidence that CPN20 is part of the CLP chaperone-protease complex assembly with a yet unknown function.

### Overlap between CLPD and CLPC1 interactors

The partial overlap in CLPD and CLPC1 interactors and overlap between CLPD trapped proteins and upregulation in *clpc1* backgrounds is easiest explained by mixed CLPC1-CLPD oligomers. Some of the proteins that were only found in the CLD traps and not in the CLPC1 traps and were not upregulated in *clpc1*, *clps1* or *clp* protease mutants are potential candidates for substrates specific to CLPD. However, follow-up experiments will be needed to validate these candidates as CLP substrates.

## Materials and Methods

### Plant growth and materials

The *Arabidopsis thaliana* (Col-0) T-DNA insertion line *clpd-1* (SAIL_77_G05; *CLPD* is AT5G51070) is described in (Nishimura et al., 2013), the *clpc1-1* null line is described in (Sjogren et al., 2004; Kovacheva et al., 2005; Nishimura et al., 2013), and the *35S*:CGEP-S718R line is described in (Bhuiyan et al., 2020).

Seeds were vernalized for 48 hours at 4°C and then grown on half-strength Murashige and Skoog (½ MS) plates with 1% sucrose and 0.6% Phytoblend at 100 µmol photons s^-1^ m^-2^ light intensity, 10 hours light/14 hours dark, 22°C day/20°C night temperature, and 60% relative humidity. After two weeks, seedlings were transferred to soil and grown in a light/dark regime of 10h/14h (short day) or 16/8 (long day) at 22°C day/20°C night temperature, 60% relative humidity, and 130 µmol photons s^-1^m^-2^.

For fresh weight measurement and chlorophyll determination, seedlings were grown on plates as described. Whole seedlings with the root removed were collected, weighed immediately, and flash frozen in liquid nitrogen for pigment isolation.

For comparison of *CLPD* mRNA and protein accumulation levels throughout different developmental stages, wild type Col-0 *Arabidopsis* seeds were grown as described in long day conditions after transfer to soil. The stage 1.02 (defined as having two true leaves over one mm in length) sample was collected from seedlings growing on ½ MS plates but all other samples were taken after transfer to soil. For stage 1.02, 1.05 (5 rosette leaves over one mm in length), and 1.14 (14 rosette leaves over one mm in length) samples the entire above ground seedling (excluding the roots) was collected, flash frozen in liquid N_2_ for subsequent RNA extraction. For the stage 3.7 (rosette is 70% of final size), 5.10 (first flower buds are visible), 6.30 (30% of flowers to be produced have opened), and 6.75 (75% of flowers to be produced have opened) samples, the entire rosette was ground in order to eliminate the variability likely to result from trying to identify leaf number in very old plants. Portions of each ground rosette were set aside for RNA and protein extraction so that RNA and protein levels could be compared within the same tissue sample. Three biological replicates were collected for every developmental stage (one individual plant per replicate), and all samples were harvested approximately two hours after the beginning of the light period.

### Pigment determination

For chlorophyll determination, individual seedlings were ground to a fine powder in liquid nitrogen and 500 µL ice cold 80% acetone was added to each. The samples were vortexed and incubated overnight at 4°C in the dark then spun at 10,000x*g* for five minutes at 4°C. The supernatant was added to 500 µl ice cold 80% acetone and absorbance was read at 663.2, 646.8 and 470nm. Chlorophyll a,b and carotenoid concentration were calculated as in (Lichtenthaler, 1987).

### RNA extraction, cDNA synthesis, and qRT-PCR

RNA extraction was performed with the Qiagen RNeasy Mini Kit with on-column DNAse digestion. cDNA synthesis was carried out with the Invitrogen SuperScript III First-Strand Synthesis Kit. For measuring *CLPD* mRNA levels through development, qRT-PCR was performed with PowerTrack SYBR Green Master Mix with three technical replicates per reaction. For measuring transgene expression in T_3_ *35S*:CLPD-STREP plants, qRT-PCR was performed with iTaq Universal SYBR Green Supermix and the reverse primer contained sequence in the *STREPII* tag to prevent cross-amplification of endogenous *CLPD*. All qRT-PCR experiments were conducted with a QuantStudio 7 Pro qPCR machine and results were normalized to the geometric mean of *ACTIN2* (*ACT2* – AT3G18780) and *UBIQUITIN10* (*UBQ10* – AT4G05320) levels. All primer sequences can be found in Supplemental Table S2.

### Generating transgenic lines

We generated transgenic lines of CLPD-STREPII and CLPD-TRAP-STREPII in the wild type *Arabidopsis thaliana* (Col-0) background that differed in their promoters: *35S* constitutive promotor (BASTA) or genomic *CLPD* (*gCLPD*) promoter (hygromycin). The *CLPD-WT* sequence was amplified from a cDNA library and the sequence coding for a C-terminal *STREPII*-tag was introduced by the reverse primer at the 3’ end. The Walker B domain E388A and E737A mutations were introduced in the coding sequence of *CLPD* by overlap extension PCR. Both *CLPD-STREP* and *CLPD-TRAP-STREP* were cloned into the gateway entry vector pCR8-GW (Invitrogen) and subsequently ligated into binary vector pEARLYGATE100 by LR reaction. pEARLYGATE100 contains a *1x35S* promoter and BASTA as selection marker for plant transformation. For expression of CLPD-STREP and CLPD-TRAP-STREP with the genomic *CLPD* promoter, 888 bp of the 5’ region upstream of the *CLPD* gene was amplified from genomic DNA and ligated into the pMDC201 vector at the HindIII and XbaI sites to replace the *2x35S* promoter on the pMDC201 vector backbone. The pMDC201 vector carries a hygromycin selection marker. The *CLPD* coding sequence was added to the plasmid from the gateway entry vector by LR reaction. The direction of the insertion was further confirmed by PCR and DNA sequencing. All primers can be found in Supplemental Table S2.

All plasmids were transformed into *Agrobacterium tumefaciens* G3101 strain by electroporation following the electroporation device manufacturer’s instruction (BioRad). The positive clones were selected on rifampicin (10 µg/ml) and gentamycin (50 µg/ml) (for the agrobacterial strain) and kanamycin (50 µg/ml) (for the plasmid). Wild type Col-0 Arabidopsis plants were transformed with agrobacteria by the floral dip method (Zhang et al., 2006). Positive transgenics were selected by growing seeds either on glufosinate ammonium (BASTA) (12.5 µg/mL) (*35S* lines) or hygromycin (15 µg/ml) (*gCLPD* lines) MS plates. Insertion of transgenes was verified by PCR-based genotyping and accumulation of transgenic protein was confirmed by immunoblot (anti-STREPII). Presence or absence of the trapping E388A and E737A mutations in the Walker B domains was confirmed by DNA sequencing.

### Protein extraction for MSMS and immunoblotting analyses

For extraction of soluble cellular proteins for MSMS analysis of chaperone accumulation levels through development and for measuring transgenic protein accumulation levels in *35S*:CLPD-STREP plants by immunoblotting, tissue was ground in liquid nitrogen into a fine powder. One mL of wash buffer per gram of tissue was added (wash buffer: 50mM HEPES-KOH, 10mM MgCl_2_, 75mM NaCl, 15% glycerol pH 7.8), and samples were vortexed three times for 30 seconds each. Samples were then spun at 10,000x*g* for 15 minutes and the supernatant was collected and used immediately for SDS-PAGE or flash frozen in liquid nitrogen.

For the ‘before bolting’ (stage 3.50) affinity CLPD trapping experiment, *35S*:CLPD-TRAP-STREP and *35S*:CGEP-STREP plants were grown on soil as described above in short day conditions and tissue was harvested when the plants reached developmental stage 3.50. The light period of the day prior to harvesting was shortened by two hours. One hour after the light period began the next morning, whole rosettes (detached from the roots) from six to eight individual plants were harvested and pooled into samples of five grams each. Tissue collection and protein extraction from both *35S*:CLPD-TRAP-STREP and *35S*:CGEP -STREP plants were carried out three times on three different days to generate the three biological replicates per genotype. All subsequent steps were carried out at 4°C in the dark under green light. For crude chloroplast isolation, fresh tissue was blended in 100mL ice-cold grinding buffer three times for ten seconds each (grinding buffer: 50mM HEPES-KOH, 330mM sorbitol, 2mM EDTA-Na_2_, 5mM ascorbic acid, 5mM cysteine, pH 8). The blended sample was filtered through two layers of Miracloth with 22µm pore size. Chloroplasts were collected by spinning for five minutes at 1300x*g*. The crude chloroplast pellet was resuspended in two mL resuspension buffer and incubated for 15 minutes on ice (resuspension buffer: 50mM HEPES-KOH, 10mM MgCl_2_, 75mM NaCl, 250µg/µL Pefabloc, 250µg/µL Avidin, 5mM cysteine, 5mM ascorbic acid, 15% glycerol, pH 7.8). Samples were vortexed three times for 30 seconds each and then spun at 100,000x*g* for 1.5 hours. The supernatant was collected as soluble crude chloroplast protein extract and used immediately for affinity purification.

For the ‘after bolting’ (stage 6.00) trapping experiment, *35S*:CLPD-TRAP-STREP and *35S*:CGEP-STREP plants were grown on soil as described above in long day conditions and tissue was harvested the day before reaching growth stage 6.00 (first flower buds opening). Tissue was harvested around one hour after the beginning of the light period, and the light period of the previous day was shortened by two hours. The whole rosettes (detached from bolts and roots) from around six individual plants were harvested and pooled to make 10-gram samples (three samples per genotype were harvested as biological replicates) and flash frozen in liquid nitrogen. To extract total soluble protein, each leaf tissue sample was finely ground in liquid N_2_ using a mortar and pestle, then added to one mL of extraction buffer per gram of tissue (extraction buffer: 50mM HEPES-KOH, 15% glycerol, 10mM MgCl_2_, 75mM NaCl, 5mM ascorbic acid, 250 µg/mL Pefabloc serine protease inhibitor, and 250 µg/mL avidin, pH 7.8). The samples were vortexed three times for thirty seconds each and filtered through four layers of 20μm Miracloth, then spun at 100,000x*g* for 1.5 hours. The soluble protein supernatant was collected and used directly for affinity purification. All protein extraction steps were carried out in the dark under green light at 4°C.

### Affinity purification

For StrepII affinity purification, Streptactin XT 4Flow columns (IBA Life Sciences) were prepared as in (Schmidt and Skerra, 2007) with 1mL bed volume. Prior to use, columns were equilibrated in two column volumes wash buffer (wash buffer: 50mM HEPES-KOH, 10mM MgCl_2_, 75mM NaCl, 15% glycerol pH 7.8). The soluble protein extract was applied to the columns, the flow through was discarded, and the columns were washed with 10 column volumes of wash buffer. Proteins were eluted with four column volumes of elution buffer (wash buffer with 50mM biotin, pH 7.8) and concentrated in Amicon 4mL (3 kDa cutoff) ultra centrifugal filters by centrifugation at 4500x*g*. Concentrated eluates were aliquoted and stored at -80°C for further proteome and MSMS analysis. Columns were regenerated between each replicate with 15 column volumes 3M MgCl_2_, followed by 8 column volumes of storage buffer (storage buffer: 50mM HEPES-KOH, 10mM MgCl_2_, 75mM NaCl, pH 7.8). All purification and concentration steps were performed at 4°C.

### SDS-PAGE and immunoblot analysis

For immunoblot analysis, protein aliquots of either soluble leaf protein or affinity eluates were separated by SDS-PAGE using homemade 6% acrylamide stacking and 12% acrylamide separation gels, followed by transfer to 0.2 μm pore size nitrocellulose membrane, staining with Ponceau S, and detection by chemiluminescence. Antisera used were anti-CLPR2 (Asakura et al., 2012) and anti-CLPT2 (Kim et al., 2015) produced by the van Wijk lab against recombinant protein, and anti-STREPII (1:2500, Genscript). Protein concentrations were determined using the BCA Protein Assay Kit (ThermoFisher).

### Proteomics and MSMS

Protein samples were separated by SDS-PAGE on Biorad Criterion Tris-HCl precast gels (10.5-14% acrylamide gradient) with three biological replicates. For the analysis of chaperone protein accumulation levels through development, one band was cut from each gel lane using the marker as a guide to excise from just above 100 kDa to just below 75 kDa and only this gel piece was further processed. For SDS-PAGE gels containing the affinity eluates of the CLPD trapping experiments, the entirety each gel lane was used, cut into consecutive gel slices (eight slices per lane for the trapping experiment at stage 3.50 and two slices per lane for the experiment at stage 6.00). Each gel slice underwent reduction, alkylation, and in-gel digestion with trypsin as described in (Friso et al., 2011). The peptides were resuspended in 2% formic acid and analyzed using a QExactive mass spectrometer equipped with a nanospray flex ion source and interfaced with a nanoLC system and autosampler (Dionex Ultimate 3000 Binary RSLCnano system) as described in (Rei Liao et al., 2022) with the exception that AGC target values were set at 1 x 10^6^ for the MSMS survey scans and maximum scan time 30 ms and 5.10^5^ for MSMS scans and maximum scan time 50 ms.

For the developmental series on the abundance of the chaperones (CLPC1,2, CLPD, cpHSP90 and CLPB3), protein identification and label-free intensity-based quantification were conducted using the FragPipe computational platform (ver. 23) with the LFQ-MBR preset workflow with the following modifications. MSBooster in MSFragger and Match between Runs (MBR) in the Quant MS1 tab were deactivated. The peptides were searched against the Araport 11 database using MSFragger (ver. 4.2) (Kong et al., 2017; Teo et al., 2021), applying carbamidomethylation of cysteine as a fixed modification, as well as N-terminal acetylation, formylation, and methionine oxidation as variable modifications. MSFragger search results were validated with Percolator and ProteinProphet integrated into FragPipe (Nesvizhskii et al., 2003; Kall et al., 2007; da Veiga Leprevost et al., 2020). Peptide intensity was calculated using IonQuant (ver. 1.11.9) (Yu et al., 2021). The FragPipe-generated “combined_protein” CSV file was used for further data filtering and statistical analysis with Perseus (ver2.1.4.0) (Tyanova et al., 2016). First, the decoys and contaminants were manually removed from the protein table. Next, proteins with a total peptide count of fewer than two were discarded. The resulting protein file was uploaded to Perseus.

Protein identification and quantification for the two CLPD-trapping experiments (stage 3.50 and 6.00) were done using the MASCOT search engine and spectral counting for quantification, essentially as described (Rei Liao et al., 2022), with carbamidomethylation of cysteine as a fixed modification and N-terminal acetylation, formylation, and methionine oxidation as variable modifications.

## Supporting information

Suppl Data set S1

## Acknowledgments

We thank Claire Ravenburg for discussions and support.

## Author Contributions

M.Y.A, P.R., N.H.B, B.Y. and K.J.V.W. conceived and designed and carried out experiments. All authors analyzed the results. M.Y.A and K.J.V.W. wrote the article. All authors read and contributed to the final article.

## Supplemental material

The following materials are available in the online versions of this article

**Supplemental Figure S1.** Analysis of CLP mRNA and protein accumulation data mined from ePLANT, ATHENA and from (Tamary et al., 2019).

**Supplemental Figure S2.** Validation of the *clpd-1* null t-DNA insertion line.

**Supplemental Figure S3.** Analysis of T3 generation of homozygous *35S*:CLPD-STREP-1 and 2 lines at the seedling stage.

**Supplemental Figure S4.** Analysis of T2 and T3 generations of homozygous *35S:*CLPD-STREP-1 and 2 lines after 3- and 7-weeks growth on soil.

**Supplemental Figure S5.** Characterization of the *35S*:CLPD-TRAP-STREP line

**Supplemental Figure S6.** Characterization of transgenic lines expressing *gCLPD:*CLPD-TRAP-STREP and *gCLPD:*CLPD-TRAP-STREP

**Supplemental Table S1.** Top 50 co-expressors of CLPC1, CLPC2, CLPB3, and cpHSP90 and their overlap - from ATTED u4

**Supplemental Table S2.** Primers used in this study.

**Supplemental Dataset S1.** MSMS-based proteomics analysis of the affinity eluates of 35S-CLPD-TRAP and 35S-CGEP at stage 3.50 or 6.00.

## Data availability

The data generated in this study are included in this article and the online supplementary material.

