## supplemental_figures for "The chloroplast CLPD chaperone: consequences of under- and overexpression, interaction with the CLP protease core, and candidate substrates"

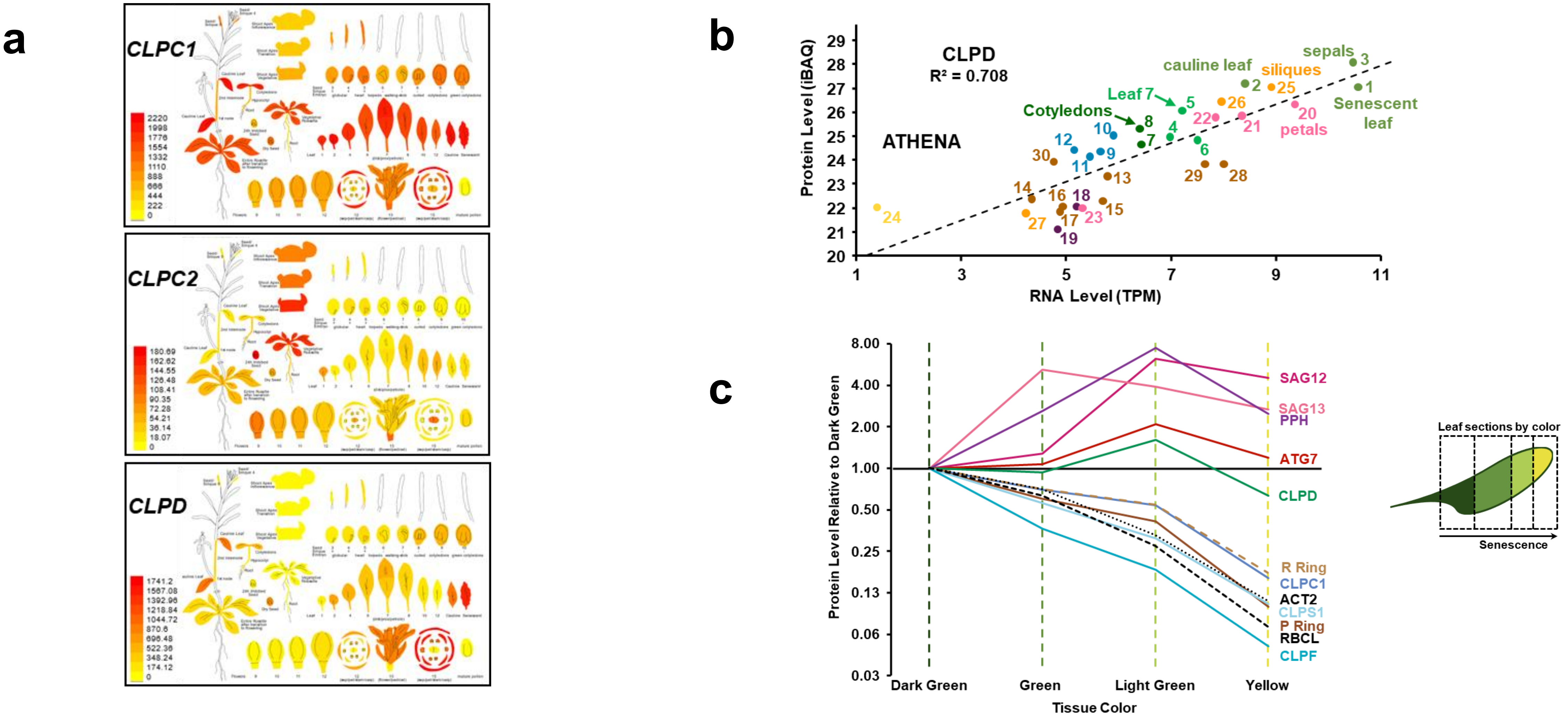


**Supplemental Figure S1**. **Supplemental Figure S1**. Analysis of CLP mRNA and protein accumulation data mined from ePLANT, ATHENA and from (Tamary et al., 2019). a) ePLANT generated heat map display of *CLPC1*, *CLPC2*, and *CLPD* mRNA expression levels in different plant tissues (Fucile et al., 2011). Each gene map is colored relative to the maximum gene expression level of that gene so that each gene is on a different, local scale. b) CLPD mRNA and protein expression in thirty tissue types was mined from the *Arabidopsis thaliana* Expression Atlas (ATHENA) database (Mergner et al., 2020) and plotted to display correlation between RNA and protein expression. Values are in units of transcripts per million (TPM) for RNA expression and Intensity-Based Absolute Quantification (iBAQ) for protein expression. A linear regression (dotted line) showed $R^2$ = 0.708. Each point represents a different tissue sampled, labeled as follows: 1) senescent leaf; 2) 1st cauline leaf; 3) sepal; 4) rosette leaf 7, petiole; 5) rosette leaf 7, proximal part; 6) rosette leaf 7, distal part; 7) shoot apical meristem, cotyledons and first leaves; 8) cotyledons; 9) stem, 1st node; 10) stem, 2nd internode; 11) hypocotyl; 12) flower pedicle; 13) root cell culture; 14) root cell culture; 15) root tip; 16) root; 17) root upper zone; 18) callus; 19) egg-cell like callus; 20) petal; 21) stamen; 22) flower; 23) carpel; 24) pollen; 25) silique valves; 26) silique septum; 27) silique (stage 3); 28) seed (embryo stage 10); 29) seed (mature, dry); 30) seed (mature, imbibed). c) Protein levels within different sectors of senescent leaves sampled by tissue color as a metric for progressing senescence: dark green, green, light green, and yellow (ordered from earliest to latest in senescence). Accumulation levels of CLP proteins, proteins involved in leaf senescence, the Rubisco large subunit (RBCL), and ACTIN2 (ACT2) presented relative to the amount of each individual protein in dark green tissue (Tamary et al., 2019).

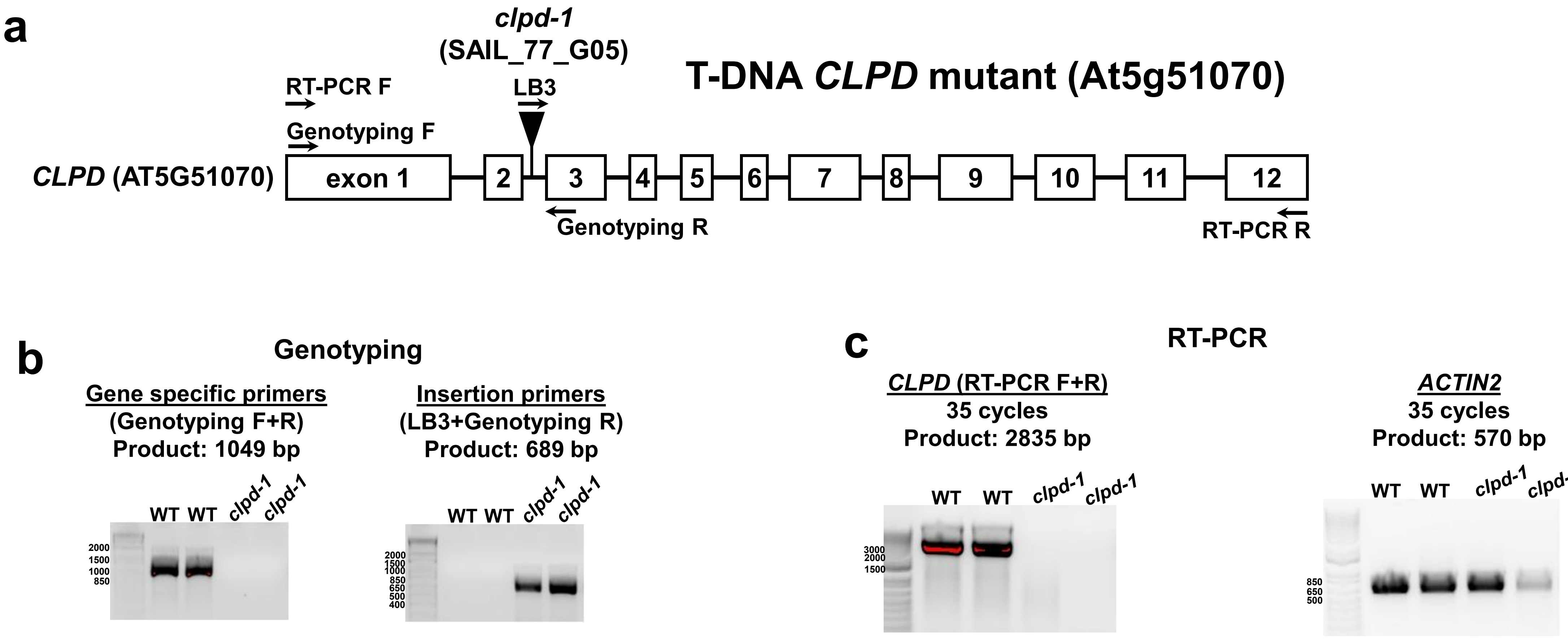


**Supplemental Figure S2**. Validation of the *clpd-1* null t-DNA insertion line. **a)** Diagram of the *CLPD* gene with boxes representing exons, connecting lines representing introns, arrows representing primer locations, and the black triangle indicating the position of the *clpd-1* (SAIL_77_G05) t-DNA insertion. **b)** PCR of gDNA extracted from wild type and *clpd-1* plants with primers specific to the endogenous *CLPD* gene (left) and primers for the t-DNA insertion (right). Homozygosity of the *clpd-1* line is demonstrated by absence of amplification in the gene-specific PCR and presence of amplification in the insertion PCR. **c)** RT-PCR of cDNA synthesized from RNA extracted from wild type and *clpd-1* plants. RT-PCR was performed with primers for endogenous *CLPD* (to verify disrupted *CLPD* expression in the *clpd-1* null line) and *ACTIN2* as a housekeeping gene. All reactions were done with 35 cycles.

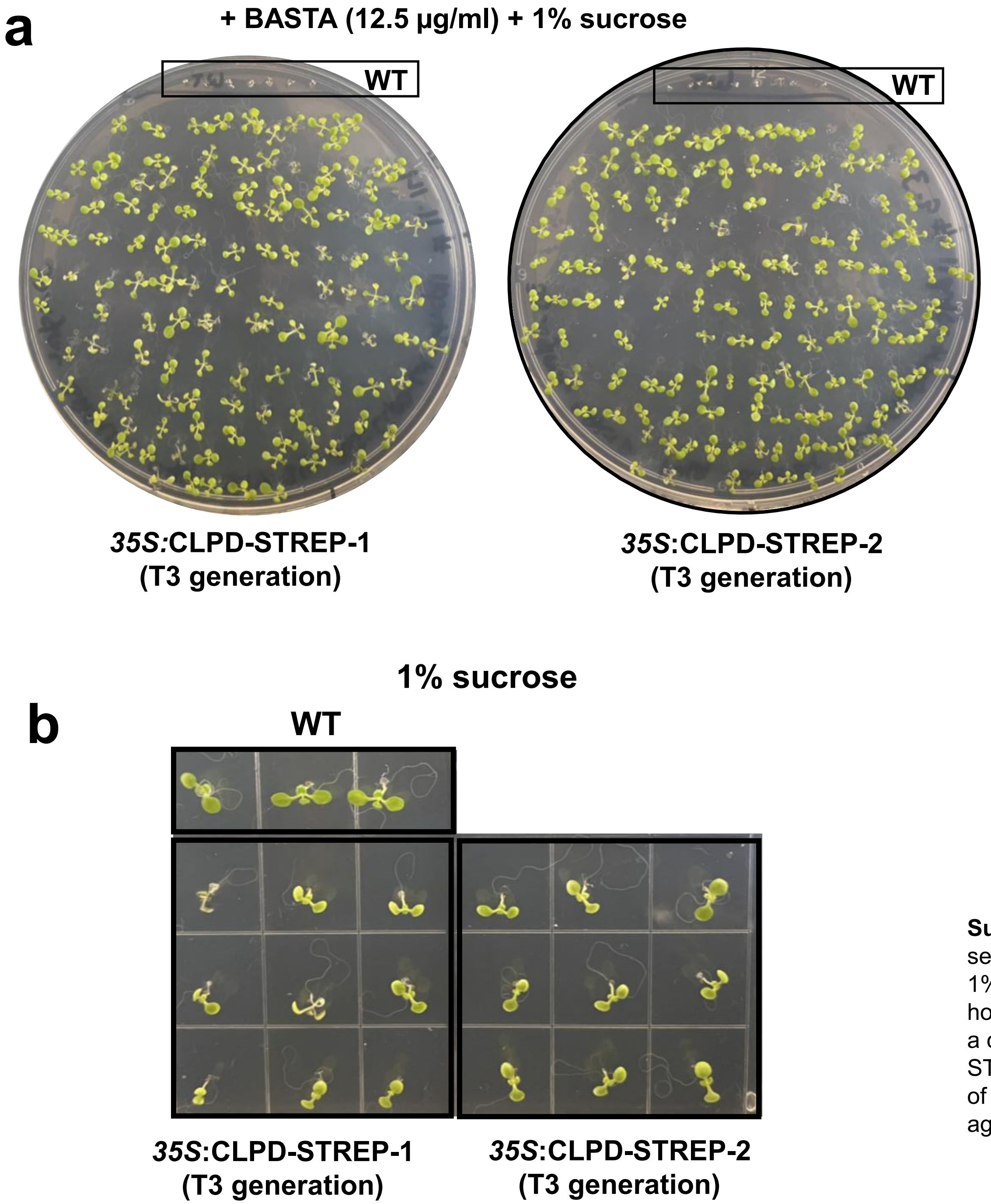


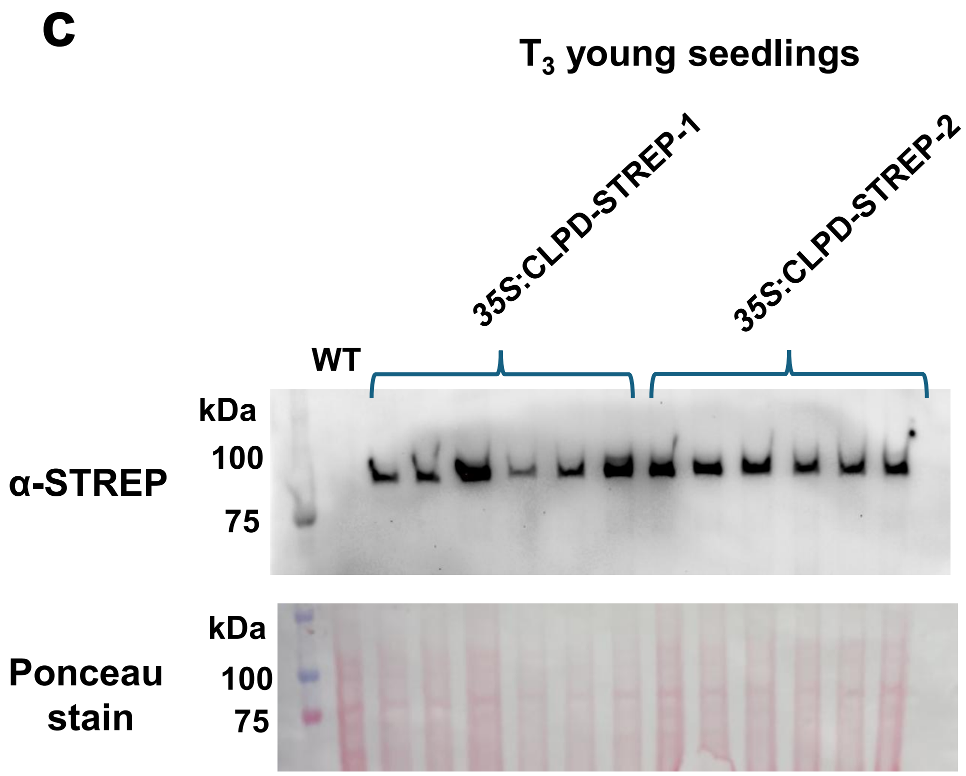


**Supplemental Figure S3.** Analysis of T3 generation of homozygous *35S*:CLPD-STREP-1 and 2 lines at the seedling stage. a) $T_3$ *35S:*CLPD-STREP-1 and *35S:*CLPD-STREP-2 seedlings grown on ½ MS plates with 1% sucrose and 12.5 µg/mL glufosinate ammonium (BASTA) for selection of the transgene to verify homozygosity of both lines. A few wild type seeds were grown on the same plates (in boxes on top) to act as a control to demonstrate the phenotype of BASTA susceptibility. **b)** $T_3$ *35S:*CLPD-STREP-1 and *35S:*CLPD-STREP-2 seedlings grown on ½ MS plates with 1% sucrose (and no selection). **c)** Anti-STREPII immunoblot of protein extracted from individual $T_3$ *35S:*CLPD-STREP-1 and *35S:*CLPD-STREP-2 seedlings at the same age and grown in the same conditions as those used for phenotypic analysis in (Figure 5).

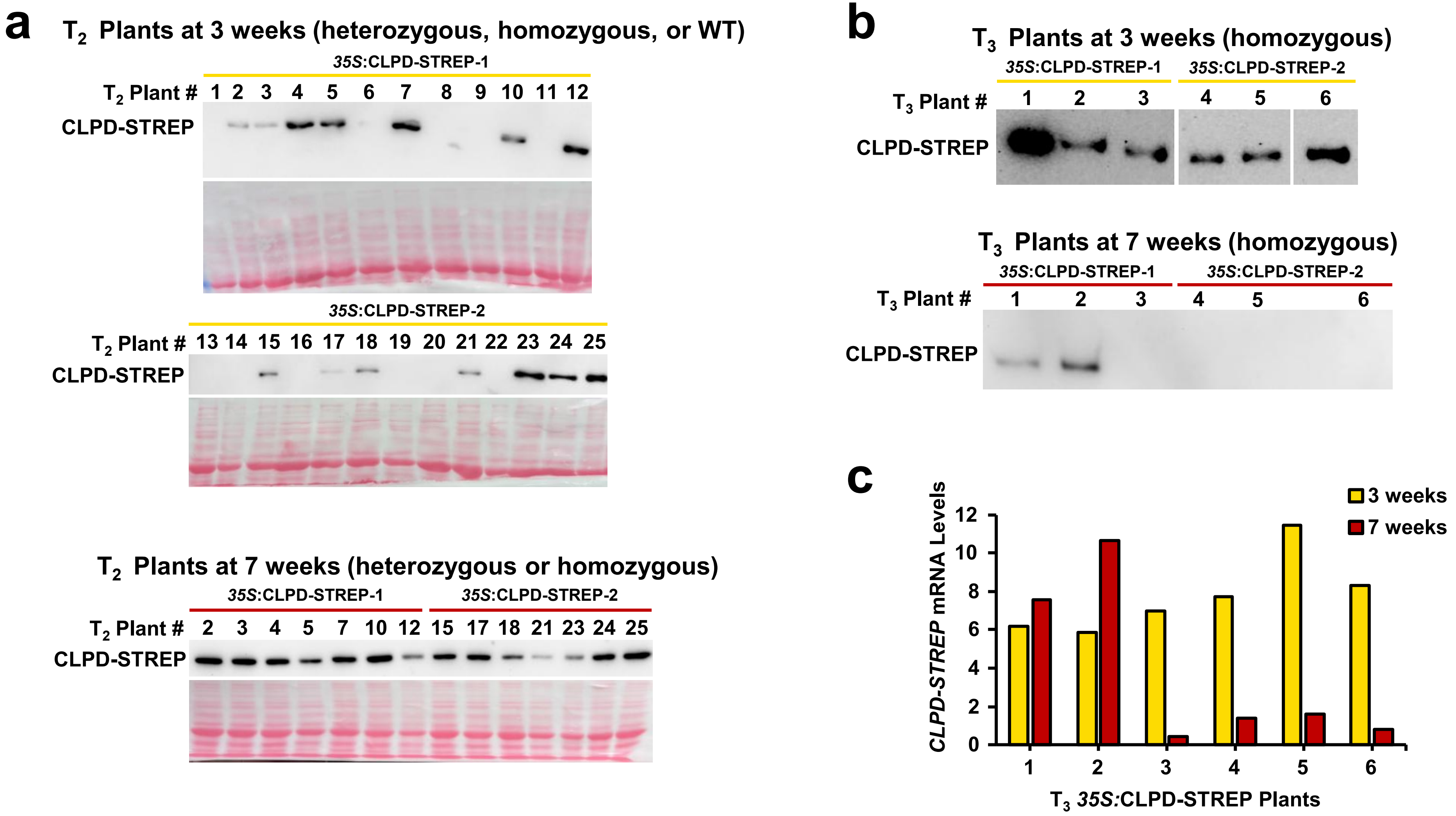


**Supplemental Figure S4.** Analysis of T2 and T3 generations of homozygous *35S:*CLPD-STREP-1 and 2 lines after 3- and 7-weeks growth on soil. a) Anti-STREPII immunoblot of leaves from a segregating population (no antibiotic selection was used) of 26 $T_2$ *35S:*CLPD-STREP plants (WT, heterozygous or homozygous for the transgenes) at three weeks and seven weeks. Every $T_2$ plant that had protein expression at three weeks (heterozygous or homozygous for the transgene) maintained protein expression at seven weeks. Plants #1-12 are from the *35:*CLPD-STREP-1 line and plants 13-25 are from the *35:*CLPD-STREP-2 line. b) Anti-STREPII immunoblot of protein extracted from leaves from homozygous $T_3$ *35:*CLPD-STREP-1,2 plants at three weeks and seven weeks. Plants #1-3 are from the *35:*CLPD-STREP-2 line and plants #4-6 are from the *35:*CLPD-STREP-1 line. c) RNA was extracted from leaves of the same plants as used in B and qRT-PCR was performed with primers specific for transgenic *CLPD-STREP* transcripts, showing that loss of transgene expression occurs on the mRNA level.

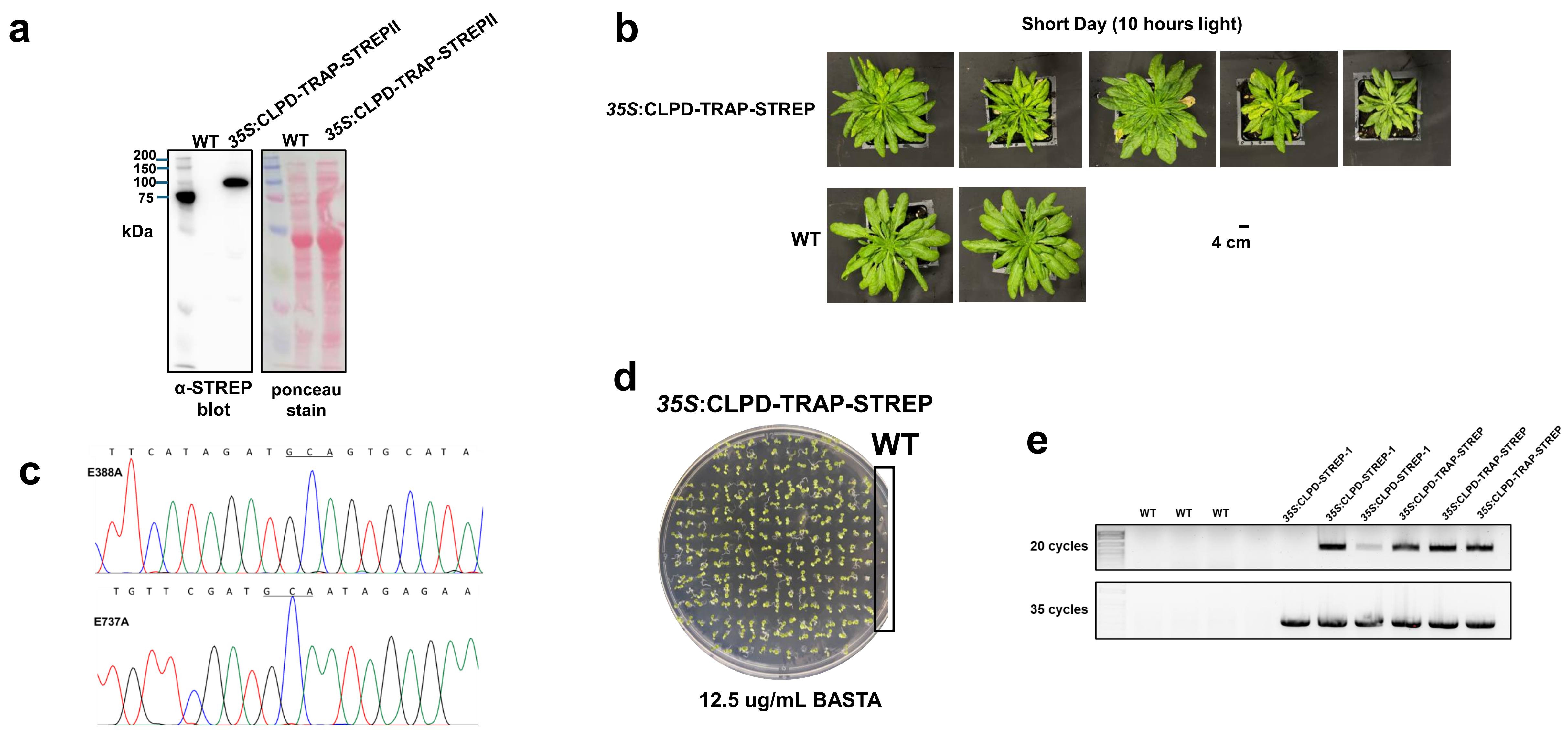


**Supplemental Figure S5.** Characterization of the *35S*:CLPD-TRAP-STREP line. **a)** SDS-PAGE and immunoblotting with anti-STREPII serum of total protein extracted from wild type and *35S:*CLPD-TRAP-STREP plants. **b)** *35S*:CLPD-TRAP-STREP phenotype grown in short day condition in stage 3.75. Pictures of *35S*:CLPD-TRAP-STREP and wild type plants at ~stage 3.75 (shortly before bolting) grown in short day conditions. All *35S*:CLPD-TRAP-STREP plants are the progeny of the same homozygous parent. **c)** Sanger sequencing of both Walker B domains of amplified transgenic CLPD DNA sequence from a *35S:*CLPD-TRAP-STREP plant shows presence of both trap mutations, seen as alanine (GCA). d**)** *35S:*CLPD-TRAP-STREP seedlings grown on ½ MS plates with 1% sucrose and 12.5 µg/mL glufosinate ammonium (BASTA) to select for presence of the transgene and verify homozygosity of the line. Several wild type seedings were grown on the same plate (in box on right) to demonstrate the phenotype of BASTA susceptibility. **e**) RT-PCR for CLPD transgenes of cDNA synthesized from RNA extracted from 35S:CLPD-TRAP-STREP and WT plants using primers for the transgene to demonstrate expression of the transgene at the mRNA level. In addition, three lanes for 35S:CLPD-WT-STREP-1 lines are shown with two of them displaying silencing.

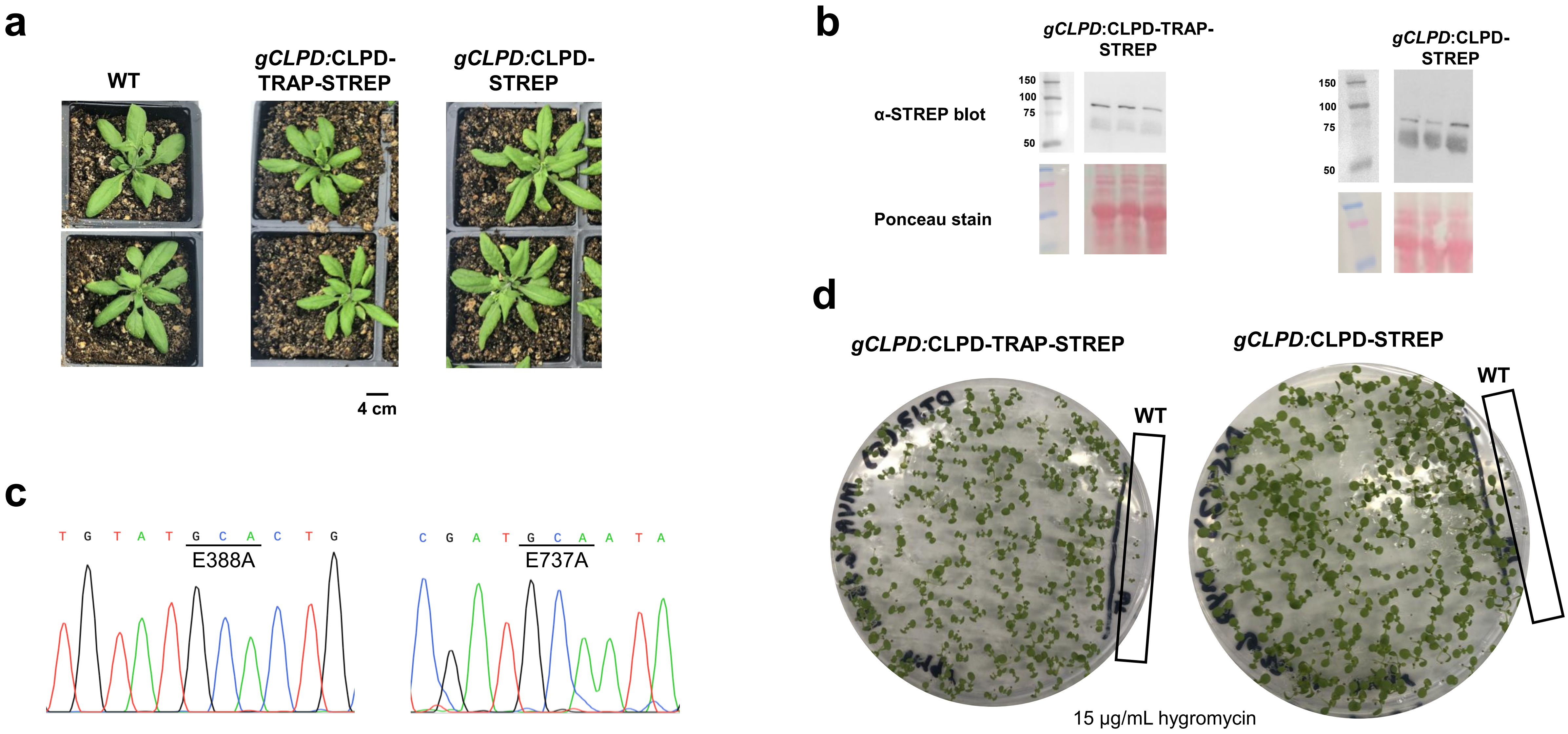


**Supplemental Figure S6.** Characterization of transgenic lines expressing *gCLPD:*CLPD-TRAP-STREP and *gCLPD:*CLPD-TRAP-STREP **a)** *gCLPD:*CLPD-TRAP-STREP, *gCLPD:*CLPD-STREP, and wild type plants grown in long day (16 hours light) conditions, shortly before bolting. **b)** Anti-STREPII immunoblot of total protein samples extracted from three *gCLPD:*CLPD-TRAP-STREP and *gCLPD:*CLPD-STREP plants. **c)** Sanger sequencing of amplified transgenic *CLPD* DNA sequence from a *gCLPD:*CLPD-TRAP-STREP plant at both Walker B domains showing the trap mutations E388A and E737A (GCA, alanine). **d)** *gCLPD:*CLPD-TRAP-STREP and *gCLPD:*CLPD-STREP seedlings grown on plates with 15 µg/mL hygromycin to select for presence of the transgene and verify homozygosity of each line. A few wild type seedlings (without hygromycin resistance) were also grown on each plate for phenotypic comparison (in box on right).
