## supplemental_text_tables for "The chloroplast CLPD chaperone: consequences of under- and overexpression, interaction with the CLP protease core, and candidate substrates"

**Supplemental Information to Annis et al.**

**Supplemental Text**

**Results**

*Linear, positive correlation between CLPD mRNA and protein levels across diverse Arabidopsis tissues and organs.* Earlier reports suggested that although CLPD mRNA increases during senescence and drought, the CLPD protein level does not (Weaver et al., 1999). To try and clarify this contradiction from data in the public domain, we mined transcript and protein abundance data for CLPD across 30 tissue types from the *Arabidopsis* Expression Atlas (ATHENA) database (there are no data for CLPC2) (Mergner et al., 2020). Supplemental Figure S1b shows a positive linear correlation between mRNA and protein for both CLPC1 and CLPD across these diverse tissues ($R^2$ = 0.609 and 0.708, respectively). The highest CLPD protein and mRNA levels were found in senescent leaves and sepals, followed by petals and siliques which also have features of senescent tissue.

*CLPD protein accumulation patterns during leaf senescence compared to CLPC1 and the CLP protease subunits.* To determine CLPD protein levels during leaf development and senescence, and to compare CLPD protein levels with CLPC1 and the rest of the CLP machinery (protease subunits and adaptors), tandem mass spectrometry (MSMS) data were extracted from an excellent study on the impact of leaf senescence on the Arabidopsis proteome (Tamary et al., 2019) (Supplemental Figure S1c). This study compared leaf proteomes during the progression of senescence, sampled by color of tissue within senescent rosettes (dark green, green, light green, and yellow leaf sections) rather than between plants of different developmental stages (Tamary et al., 2019). We re-analyzed the data to display the dynamics of the chaperones CLPD and CLPC1, the CLP core subunits of the P ring (CLPP3-6) and the R ring (CLPP1,R1-4), the accessory proteins CLPT1 and CLPT2, and the adaptors CLPS1 and CLPF. Abundances of all chloroplast CLP proteins continually decrease as senescence progresses, except for CLPD, which is highest in light green tissue (Supplemental Figure S1c). This is a pattern mirrored by several marker proteins for senescence, including SENESCENCE-ASSOCIATED GENE 12 (SAG12), SENESCENCE-ASSOCIATED GENE 13 (SAG13), PHEOPHYTIN PHEOPHORBIDE HYDROLASE (PPH), and AUTOPHAGY-RELATED GENE 7 (ATG7) (Supplemental Figure 1).

**Cited references**

**Fucile G, Di Biase D, Nahal H, La G, Khodabandeh S, Chen Y, Easley K, Christendat D, Kelley L, Provart NJ** (2011) ePlant and the 3D data display initiative: integrative systems biology on the world wide web. PLoS One **6:** e15237

**Mergner J, Frejno M, List M, Papacek M, Chen X, Chaudhary A, Samaras P, Richter S, Shikata H, Messerer M, Lang D, Altmann S, Cyprys P, Zolg DP, Mathieson T, Bantscheff M, Hazarika RR, Schmidt T, Dawid C, Dunkel A, Hofmann T, Sprunck S, Falter-Braun P, Johannes F, Mayer KFX, Jurgens G, Wilhelm M, Baumbach J, Grill**

**E, Schneitz K, Schwechheimer C, Kuster B** (2020) Mass-spectrometry-based draft of the Arabidopsis proteome. Nature **579:** 409-414

**Rei Liao JY, Friso G, Forsythe ES, Michel EJS, Williams AM, Boguraev SS, Ponnala L, Sloan DB, van Wijk KJ** (2022) Proteomics, phylogenetics, and coexpression analyses indicate novel interactions in the plastid CLP chaperone-protease system. J Biol Chem **298:** 101609

**Tamary E, Nevo R, Naveh L, Levin-Zaidman S, Kiss V, Savidor A, Levin Y, Eyal Y, Reich Z, Adam Z** (2019) Chlorophyll catabolism precedes changes in chloroplast structure and proteome during leaf senescence. Plant Direct **3:** e00127

**Weaver LM, Froehlich JE, Amasino RM** (1999) Chloroplast-targeted ERD1 protein declines but its mRNA increases during senescence in Arabidopsis. Plant Physiol **119:** 1209-1216

**Supplemental Table S1. Top 50 co-expressors of CLPC1, CLPC2, CLPB3, and cpHSP90 and their overlap - from ATTED u4**

| ath-u.4 ClpD | Accession | Gene | ath-u.4 / CLPC1 | Accession | Gene | ath-u.4 ClpC2 | Accession | Gene | ath-u.4 / CLPB3 | Accession | Gene | ath-u.4 /cpHSP90 | Accession | Gene |
|---|---|---|---|---|---|---|---|---|---|---|---|---|---|---|
| 9.7 | AT1G13990.1 | AT1G13990 | 10.2 | AT2G30950.1 | VAR2 | 6.6 | AT3G46740.1 | TOC75-III | 8.9 | AT3G11670.1 | DGD1 | 15.6 | AT2G28000.1 | CPN60A |
| 9.2 | AT5G13800.1 | PPH | 8.8 | AT3G19170.1 | PREP1 | 5.9 | AT1G06950.1 | TIC110 | 6.8 | AT5G03650.1 | SBE2.2 | 12.3 | AT3G13470.1 | Cpn60beta2 |
| 7.3 | AT5G47860.1 | AT5G47860 | 8.7 | AT5G60600.1 | HDS | 5.7 | AT5G18960.1 | FRS12 | 6.7 | AT1G23180.1 | AT1G23180 | 11.8 | AT4G24280.1 | cpHsc70-1 |
| 7 | AT3G07700.1 | AT3G07700 | 8.6 | AT5G42270.1 | VAR1 | 5.6 | AT3G13470.1 | Cpn60beta2 | 6.3 | AT3G04870.1 | ZDS | 11.5 | AT1G59990.1 | EMB3108 |
| 6.9 | AT3G44880.1 | ACD1 | 8.2 | AT1G49970.1 | CLPR1 | 5.3 | AT2G28000.1 | CPN60A | 6 | AT5G67030.1 | ABA1 | 11.2 | AT3G07770.1 | Hsp89.1 |
| 6.8 | AT1G22930.1 | AT1G22930 | 8.1 | AT1G67280.1 | AT1G67280 | 5.2 | AT4G24280.1 | cpHsc70-1 | 6 | AT4G24280.1 | cpHsc70-1 | 10.9 | AT1G69200.1 | FLN2 |
| 6.5 | AT1G54130.1 | RSH3 | 7.2 | AT2G28800.1 | ALB3 | 5.1 | AT3G53580.1 | AT3G53580 | 6 | AT5G64840.1 | ABCF5 | 10.4 | AT3G54090.1 | FLN1 |
| 5.8 | AT3G51130.1 | AT3G51130 | 7.2 | AT4G01800.1 | AGY1 | 5.1 | AT1G79560.1 | FTSH12 | 5.9 | AT2G22240.1 | MIPS2 | 9.9 | AT5G20720.1 | CPN20 |
| 5.6 | AT4G22920.1 | NYE1 | 7.1 | AT1G62750.1 | SCO1 | 5.1 | AT4G31210.1 | AT4G31210 | 5.8 | AT4G04020.1 | FIB | 9.8 | AT5G48470.1 | AT5G48470 |
| 5.4 | AT2G24860.1 | AT2G24860 | 6.9 | AT3G04870.1 | ZDS | 5 | AT1G09620.1 | AT1G09620 | 5.7 | AT2G37240.1 | AT2G37240 | 9.7 | AT4G02990.1 | BSM |
| 5.4 | AT1G07040.1 | AT1G07040 | 6.9 | AT1G06430.1 | FTSH8 | 5 | AT5G18570.1 | EMB269 | 5.7 | AT1G32900.1 | GBSS1 | 9.5 | AT5G49910.1 | cpHsc70-2 |
| 5.4 | AT3G03470.1 | CYP89A9 | 6.9 | AT1G36320.1 | AT1G36320 | 4.9 | AT1G17850.1 | AT1G17850 | 5.6 | AT5G09590.1 | MTHSC70-2 | 8.9 | AT3G13860.1 | HSP60-3A |
| 5.2 | AT5G54080.1 | HGO | 6.7 | AT1G79600.1 | AT1G79600 | 4.9 | AT3G01370.1 | CFM2 | 5.5 | AT1G67440.1 | emb1688 | 8.8 | AT2G04530.1 | CPZ |
| 5.1 | AT3G10420.1 | SPD1 | 6.5 | AT4G34350.1 | HDR | 4.9 | AT2G44640.1 | AT2G44640 | 5.4 | AT4G13550.1 | AT4G13550 | 8.8 | AT5G64580.1 | EMB3144 |
| 5 | AT5G27280.1 | AT5G27280 | 6.5 | AT4G29060.1 | emb2726 | 4.8 | AT4G21770.1 | AT4G21770 | 5.4 | AT1G06430.1 | FTSH8 | 8.6 | AT3G23990.1 | HSP60 |
| 5 | AT1G06430.1 | FTSH8 | 6.5 | AT2G26670.1 | TED4 | 4.8 | AT1G06220.1 | MEE5 | 5.4 | AT2G35840.1 | AT2G35840 | 8.5 | AT1G79560.1 | FTSH12 |
| 4.9 | AT1G32230.1 | RCD1 | 6.4 | AT1G09130.1 | AT1G09130 | 4.8 | AT5G64960.1 | CDKC2 | 5.4 | AT1G62740.1 | Hop2 | 8.5 | AT1G06950.1 | TIC110 |
| 4.8 | AT5G11840.1 | AT5G11840 | 6.3 | AT5G42390.1 | SPP | 4.7 | AT3G04340.1 | emb2458 | 5.4 | AT5G48300.1 | ADG1 | 8.5 | AT2G31890.1 | RAP |
| 4.7 | AT2G20920.1 | AT2G20920 | 6.3 | AT2G35490.1 | AT2G35490 | 4.7 | AT1G74260.1 | PUR4 | 5.3 | AT2G04030.1 | CR88 | 8.3 | AT1G30680.1 | AT1G30680 |
| 4.6 | AT4G03030.1 | AT4G03030 | 6 | AT3G63520.1 | CCD1 | 4.7 | AT3G57610.1 | ADSS | 5.3 | AT3G24190.1 | AT3G24190 | 8.1 | AT4G26780.1 | AR192 |
| 4.5 | AT4G15530.1 | PPDK | 6 | AT1G48450.1 | AT1G48450 | 4.7 | AT1G62390.1 | Phox2 | 5.3 | AT5G43850.1 | ARD4 | 8.1 | AT4G01037.1 | WTF1 |
| 4.5 | AT4G22240.1 | AT4G22240 | 5.8 | AT5G46420.1 | AT5G46420 | 4.6 | AT5G08610.1 | PDE340 | 5.3 | AT2G25140.1 | CLPB4 | 8.1 | AT3G48500.1 | PDE312 |
| 4.5 | AT4G34890.1 | XDH1 | 5.7 | AT4G20850.1 | TPP2 | 4.5 | AT3G04260.1 | PTAC3 | 5.3 | AT1G17050.1 | SPS2 | 8 | AT1G62390.1 | Phox2 |
| 4.4 | AT4G31860.1 | AT4G31860 | 5.7 | AT5G64840.1 | ABCF5 | 4.5 | AT5G49430.1 | AT5G49430 | 5.2 | AT1G23740.1 | AOR | 8 | AT2G31840.1 | MRL7-L |
| 4.4 | AT3G14050.1 | RSH2 | 5.6 | AT1G02560.1 | CLPP5 | 4.4 | AT3G10690.1 | GYRA | 5.1 | AT4G00490.1 | BAM2 | 7.9 | AT5G08610.1 | PDE340 |
| 4.4 | AT4G27990.1 | YLMG1-2 | 5.6 | AT1G70070.1 | EMB25 | 4.4 | AT1G14810.1 | AT1G14810 | 5.1 | AT1G62750.1 | SCO1 | 7.8 | AT4G09730.1 | RH39 |
| 4.3 | AT1G53570.1 | MAP3KA | 5.6 | AT4G25650.1 | ACD1-LIKE | 4.4 | AT3G45890.1 | RUS1 | 5.1 | AT5G50920.1 | CLPC1 | 7.6 | AT1G55490.1 | CPN60B |
| 4.3 | AT4G03410.1 | AT4G03410 | 5.6 | AT1G63770.1 | AT1G63770 | 4.3 | AT5G20040.1 | IPT9 | 5.1 | AT4G39210.1 | APL3 | 7.6 | AT2G34640.1 | PTAC12 |
| 4.3 | AT1G48490.1 | AT1G48490 | 5.6 | AT5G14260.1 | AT5G14260 | 4.3 | AT2G25870.1 | AT2G25870 | 5.1 | AT1G52870.1 | AT1G52870 | 7.6 | AT4G31210.1 | AT4G31210 |
| 4.3 | AT3G27110.1 | AT3G27110 | 5.6 | AT4G14210.1 | PDS3 | 4.3 | AT3G58610.1 | AT3G58610 | 5.1 | AT1G30070.1 | AT1G30070 | 7.5 | AT5G46580.1 | AT5G46580 |
| 4.3 | AT5G18130.1 | AT5G18130 | 5.5 | AT3G18390.1 | EMB1865 | 4.3 | AT3G61780.1 | emb1703 | 5.1 | AT2G26140.1 | ftsh4 | 7.5 | AT1G48570.1 | AT1G48570 |
| 4.3 | AT5G18490.1 | AT5G18490 | 5.5 | AT1G65260.1 | PTAC4 | 4.3 | AT3G23940.1 | AT3G23940 | 5 | AT3G01180.1 | SS2 | 7.4 | AT1G23180.1 | AT1G23180 |
| 4.2 | AT1G54100.1 | ALDH7B4 | 5.5 | AT1G66670.1 | CLPP3 | 4.3 | AT2G34640.1 | PTAC12 | 5 | AT2G46915.1 | AT2G46915 | 7.4 | AT2G20020.1 | CAF1 |
| 4.2 | AT5G17380.1 | AT5G17380 | 5.4 | AT4G24280.1 | cpHsc70-1 | 4.3 | AT2G31400.1 | GUN1 | 5 | AT4G09150.1 | AT4G09150 | 7.4 | AT5G04130.1 | GYRB2 |
| 4.2 | AT4G36760.1 | APP1 | 5.4 | AT5G19540.1 | AT5G19540 | 4.2 | AT1G50200.1 | ALATS | 5 | AT1G03310.1 | DBE1 | 7.4 | AT3G10690.1 | GYRA |
| 4.2 | AT1G32790.1 | CID11 | 5.3 | AT1G08540.1 | SIG2 | 4.2 | AT1G16270.1 | AT1G16270 | 4.9 | AT1G16030.1 | Hsp70b | 7.3 | AT1G10522.1 | PRIN2 |
| 4.2 | AT4G25650.1 | ACD1-LIKE | 5.3 | AT1G50250.1 | FTSH1 | 4.2 | AT2G47940.1 | DEG2 | 4.8 | AT5G64170.1 | AT5G64170 | 7.3 | AT5G48030.1 | GFA2 |
| 4.2 | AT1G22400.1 | UGT85A1 | 5.3 | AT5G67030.1 | ABA1 | 4.2 | AT1G16350.1 | AT1G16350 | 4.8 | AT3G54660.1 | GR | 7.3 | AT1G03860.1 | PHB2 |
| 4.1 | AT5G10650.1 | AT5G10650 | 5.3 | AT5G13650.1 | SVR3 | 4.2 | AT2G32730.1 | AT2G32730 | 4.7 | AT4G14070.1 | AAE15 | 7.2 | AT2G14880.1 | AT2G14880 |
| 4.1 | AT1G27980.1 | DPL1 | 5.3 | AT4G36390.1 | AT4G36390 | 4.2 | AT3G18390.1 | EMB1865 | 4.7 | AT1G79920.1 | Hsp70-15 | 7.2 | AT5G55280.1 | FTSZ1-1 |
| 4.1 | AT3G12740.1 | ALIS1 | 5.2 | AT2G31400.1 | GUN1 | 4.2 | AT2G23350.1 | PAB4 | 4.7 | AT2G42600.1 | PPC2 | 7.1 | AT5G26860.1 | LON1 |
| 4.1 | AT2G42890.1 | ML2 | 5.2 | AT3G44880.1 | ACD1 | 4.1 | AT2G32230.1 | PRORP1 | 4.7 | AT4G14210.1 | PDS3 | 7.1 | AT1G14980.1 | CPN10 |
| 4.1 | AT5G14780.1 | FDH | 5.2 | AT4G17040.1 | CLPR4 | 4.1 | AT1G55490.1 | CPN60B | 4.7 | AT3G08590.1 | iPGAM2 | 7.1 | AT3G53560.1 | AT3G53560 |
| 4.1 | AT3G17800.1 | AT3G17800 | 5.2 | AT1G17220.1 | FUG1 | 4.1 | AT1G20960.1 | emb1507 | 4.6 | AT5G60600.1 | HDS | 7 | AT3G03630.1 | CS26 |
| 4 | AT2G20010.1 | AT2G20010 | 5.2 | AT3G27820.1 | MDAR4 | 4.1 | AT3G27740.1 | CARA | 4.6 | AT5G01410.1 | RSR4 | 7 | AT5G18820.1 | Cpn60alpha2 |
| 4 | AT1G60490.1 | VPS34 | 5.2 | AT5G44000.1 | AT5G44000 | 4 | AT1G69200.1 | FLN2 | 4.6 | AT3G01510.1 | LSF1 | 7 | AT3G14900.1 | EMB3120 |
| 4 | AT3G17000.1 | UBC32 | 5.2 | AT3G09210.1 | PTAC13 | 4 | AT4G33760.1 | AT4G33760 | 4.6 | AT5G57610.1 | AT5G57610 | 6.9 | AT2G24120.1 | SCA3 |
| 4 | AT2G37150.1 | AT2G37150 | 5.2 | AT1G70200.1 | AT1G70200 | 4 | AT5G13520.1 | AT5G13520 | 4.6 | AT2G04350.1 | LACS8 | 6.8 | AT5G09590.1 | MTHSC70-2 |
| 3.9 | AT1G04970.1 | AT1G04970 | 5.2 | AT5G03900.1 | AT5G03900 | 4 | AT5G19620.1 | OEP80 | 4.5 | AT4G35090.1 | CAT2 | 6.8 | AT5G16620.1 | TIC40 |
| 3.9 | AT4G21534.1 | SPHK2 | 5.2 | AT4G01940.1 | NFU1 | 3.9 | AT4G02510.1 | TOC159 | 4.5 | AT1G12800.1 | AT1G12800 | 6.8 | AT4G27340.1 | AT4G27340 |

- shared between CLPD & CLPC1
- shared between CLPC1 & CLPC2
- shared between CLPC1 & CLPB3
- shared between CLPC2&HSP90
- shared between CLPC1&CLPC2&CLPB3
- shared between CLPD&CLPC1&CLPB4
- shared between CLPB3&HSP90

| **Supplemental Table S2.** Primers used in this study. | | | |
|---|---|---|---|
| **Sequence** | **Description** | **primer name** | **primer #** |
| GTGACTCGACTCTGTTTAGTGCA | *UBQ10* qRT-PCR forward primer | UBQ10 | 3011 |
| CCCAACAGCTCAACACTTTCG | *UBQ10* qRT-PCR reverse primer | UBQ10 | 3012 |
| GTTTTCGTTTCTATGATGCACTTGTG | *ACT2* qRT-PCR forward primer | ACT2 | 3013 |
| GGGACTAAAACGCAAAACGAAAGC | *ACT2* qRT-PCR reverse primer | ACT2 | 3014 |
| TGCCATTGCTGAAGGACTAGCG | *CLPD* qRT-PCR forward primer | CLPD | 2992 |
| AGGGACATGATGCGTTTCGTCAAG | *CLPD* qRT-PCR reverse primer | CLPD | 2993 |
| GATGATACCGGAAACCCATCGG | *CLPD-STREP* qRT-PCR forward primer | CLPD-STREPII | 2998 |
| CTTCTCGAATTGAGGATGAGACCA | *CLPD-STREP* qRT-PCR reverse primer | CLPD-STREPII | 2942 |
| TAAAGGGAGCTTCGAGTCTC | E373A forward sequencing primer | E3737A F | 2629 |
| CAAAATCTTCACCGCATCTTC | E373A reverse sequencing primer | E3737A R | 2630 |
| CCCGACCGTCCAATTGCTGC | E388A forward sequencing primer | E388A F | 2947 |
| CGGTTCAACAACTCTGGACGG | E388A reverse sequencing primer | E388A R | 2948 |
